# Multi-level analysis of reproduction in the Antarctic midge, *Belgica antarctica*, identifies female and male accessory gland products that are altered by larval stress and impact progeny viability

**DOI:** 10.1101/796797

**Authors:** Geoffrey Finch, Sonya Nandyal, Carlie Perrieta, Benjamin Davies, Andrew J. Rosendale, Christopher J. Holmes, Josiah D. Gantz, Drew Spacht, Samuel T. Bailey, Xiaoting Chen, Kennan Oyen, Elise M. Didion, Souvik Chakraborty, Richard E. Lee, David L. Denlinger, Stephen F. Matter, Geoffrey M. Attardo, Matthew T. Weirauch, Joshua B. Benoit

**Affiliations:** Department of Biological Sciences, University of Cincinnati, Cincinnati, OH; Department of Biology, Mount St. Joseph University, Cincinnati, OH, USA; Department of Biology, Miami University, Oxford, OH; Department of Biology and Health Science, Hendrix College, Conway, AR; Departments of Entomology and Evolution, Ecology and Organismal Biology, The Ohio State University, Columbus, OH; Center for Autoimmune Genomics and Etiology, Cincinnati Children’s Hospital Medical Center, Cincinnati, OH 45229, USA; Department of Entomology and Nematology, University of California, Davis, Davis, CA 95616, USA; Divisions of Biomedical Informatics and Developmental Biology, Cincinnati Children’s Hospital Medical Center, Cincinnati, OH 45229, USA.; Department of Pediatrics, University of Cincinnati College of Medicine, Cincinnati, OH 45267, USA.

**Author notes:** Author for correspondence Joshua B. Benoit Department of Biological Sciences, University of Cincinnati, Cincinnati, OH.

**Keywords:** Antarctic midge, reproduction, stress biology, accessory glands, thermal buffering

## Abstract

The Antarctic midge, *Belgica antarctica*, is a wingless, non-biting midge endemic to Antarctica. Larval development requires at least two years, but adult life lasts only two weeks. The nonfeeding adults mate in swarms and females die shortly after oviposition. Eggs are suspended in a gel of unknown composition that is expressed from the female accessory gland. This project characterizes molecular mechanisms underlying reproduction in this midge by examining differential gene expression in whole males, females, and larvae, as well as in male and female accessory glands. Functional studies were used to assess the role of the gel encasing the eggs, as well as the impact of stress on reproductive biology. RNA-seq analyses revealed sex- and development-specific gene sets along with those associated with the accessory glands. Proteomic analyses were used to define the composition of the egg-containing gel, which is generated during multiple developmental stages and derived from both the accessory gland and other female organs. Functional studies indicate the gel provides a larval food source and thermal and dehydration buffer, all of which are critical for viability. Larval dehydration stress directly reduces production of storage proteins and key accessory gland components, a feature that impacts adult reproductive success. Modeling reveals that bouts of dehydration may significantly impact population growth. This work lays a foundation for further examination of reproduction in midges and provides new information related to general reproduction in dipterans. A key aspect is that reproduction and stress dynamics, currently understudied in polar organisms, are likely to prove critical for determining how climate change will alter survivability.

## 1 INTRODUCTION

The Antarctic midge, *Belgica antarctica,* is long-lived, wingless, and the only insect endemic to maritime Antarctica(Convey & Block, 1996; Sugg, Edwards, & Baust, 1983). It has a patchy distribution along the western coast of the Antarctic Peninsula and South Shetland Islands, where it may form large aggregations into the thousands under favorable conditions (Convey & Block, 1996; SUGG et al., 1983). The larval period lasts two years; growth and development take place during the short austral summer, and larvae overwinter encased in ice (Usher & Edwards, 1984). Larvae commonly reside in damp areas and are non-selective feeders by consuming dead plant and animal matter, algae, and microorganisms (Edwards, & Baust, 1981; Strong, 1967). Larvae are extremely tolerant of numerous stresses including cold, dehydration, and UV exposure (JBenoit, Lopez-Martinez, Elnitsky, Lee, & Denlinger, 2009; Benoit, Lopez-Martinez, Michaud, et al., 2007; Lopez-Martinez et al., 2009; Lopez-Martinez, Elnitsky, Benoit, Lee, & Denlinger, 2008; Teets et al., 2008). Adult emergence is a fairly synchronous event occurring in early summer (Edwards, & Baust, 1981), and there is some evidence for protandry (Edwards, & Baust, 1981; Harada, Lee, Denlinger, & Goto, 2014). The wingless adults mate in swarms formed on rocks and other features of the substrate (Edwards, & Baust, 1981; Harada et al., 2014). Environmental stressors in Antarctica can be severe and highly variable over short distances and time periods (Convey, 1997; Kennedy, 1993), and swarming may play a role in locating and taking advantage of intermittent, favorable microhabitats (Convey & Block, 1996; Hahn & Reinhardt, 2006; Sugg et al., 1983). Mating swarms present a potential obstacle for establishing long-lasting colonies in a laboratory setting, as mating is likely facilitated by large-scale, synchronous adult emergence (Harada et al., 2014).

Adult females that emerge in the laboratory lay, with a few exceptions, a single egg mass, each containing about 40 eggs (Convey & Block, 1996; Edwards, & Baust, 1981; Harada et al., 2014; Sugg et al., 1983). Nevertheless, multiple matings are common, and multiple oviposition events have been reported (Edwards, & Baust, 1981; Harada et al., 2014; Sugg et al., 1983). Specific underlying materials transferred from the male to the female during copulation are unknown in *B. antarctica*. Eggs are encased in a hygroscopic gel that has been suggested as a potential food source for developing larvae (Edwards, & Baust, 1981; Harada et al., 2014). This gel is likely secreted by the female accessory gland during oviposition, but the protein components of the gel are unknown. Little additional information is available on reproduction in these extremophiles (Edwards, & Baust, 1981; Harada et al., 2014), and chironomid reproduction in general is poorly studied beyond the basic descriptions of their reproductive anatomy and impaired reproduction during exposure to toxic substances (Sibley, Ankley, & Benoit, 2001; Vogt et al., 2007; Wensler & Rempel, 1962).

By contrast, reproduction in other dipteran species has been examined extensively (Alfonso-Parra et al., 2016; Avila, Sirot, LaFlamme, Rubinstein, & Wolfner, 2011; J.B. Benoit, Attardo, Baumann., Michalkova, & Aksoy, 2015; A. G. Clark et al., 2007; Dottorini et al., 2007; Izquierdo et al., 2019; K. P. Lee et al., 2008; McGraw, Clark, & Wolfner, 2008; Meier, Kotrba, & Ferrar, 1999; Papa et al., 2017; Polak et al., 2017; Ravi Ram & Wolfner, 2007; Villarreal et al., 2018). In most Diptera, insemination does not involve injection of a spermatophore; rather, male seminal fluid is usually transferred directly into the female reproductive tract, often with the addition of a mating plug to reduce male-male competition as is seen in mosquitoes and *Drosophila* (Giglioli & Mason, 1966; Lung & Wolfner, 2001; Mitchell et al., 2015; Scolari et al., 2016). Some flies do utilize a spermatophore, which may facilitate quicker mating while also creating a barrier to multiple inseminations (Kotbra, 1996). In the tsetse fly, *Glossina morsitans,* a dipteran that uses a spermatophore, proteins making up the spermatophore are secreted from the male accessory gland during copulation (Attardo et al., 2019; Scolari et al., 2016). The spermatophore is deposited such that the spermatozoa are funneled efficiently to the openings of the spermathecal ducts, allowing only one spermatophore to maintain this connection at a time (Attardo et al., 2019; Scolari et al., 2016). In mosquitoes, accessory gland-specific proteins, along with the steroid hormone, 20-hydroxy-ecdysone, are transferred to the female during copulation, producing a mating plug (Dottorini et al., 2007; Mitchell et al., 2015; Rogers et al., 2008). In *An. gambiae* the mating plug has multiple effects that promote copulation, enhance egg production, and even trigger egg laying (Dottorini et al., 2007; Gabrieli et al., 2014; Mitchell et al., 2015; Thailayil, Magnusson, Godfray, Crisanti, & Catteruccia, 2011). First-male precedence and last-male precedence have both been reported in multiple species within the Diptera (Dixon, Coyne, & Noor, 2003; Gwynne, 2012; Price, 1997; Shutt, Stables, Aboagye-Antwi, Moran, & Tripet, 2010), but it remains unknown how fertilization priority is established in *B. antarctica*. Depending on the mating strategy of the species, contents of the spermatophore may include a large amount of seminal fluid proteins (SFPs) and other factors or contain primarily the sperm itself (Avila et al., 2011; Lung & Wolfner, 2001; Rogers et al., 2008; Scolari et al., 2016). These seminal fluid proteins are suspected or demonstrated to have a variety of functions, including induction of refractoriness in the female, counteracting protease activity of female secretions with protease inhibitors, defending spermatozoa against microbial assault, and neutralizing reactive oxidative species (Alfonso-Parra et al., 2016; Avila et al., 2011; Lung & Wolfner, 2001; Ravi Ram & Wolfner, 2007; Shutt et al., 2010), all which can compromise sperm function and impair their ability to fertilize. The particular cocktail and amounts of SFPs utilized reflects the selection pressure from life-history differences, conspecific competition, and diverse reproductive strategies between species and even within species (Hopkins et al., 2019; Hopkins, Sepil, & Wigby, 2017; Izquierdo et al., 2019; Papa et al., 2017).

Secretions from the female accessory glands also play important roles in insemination and oviposition in dipterans, as well as other insects. In the house fly, *Musca domestica,* female accessory gland secretions enhance sperm viability (Degrugillier, 1985; Leopold & Degrugillier, 1973). In some species, substances secreted from the female accessory gland are expelled with the eggs at oviposition and often function as an adhesive, anchoring eggs to the substrate (Lococo & Huebner, 1980). Accessory gland secretions may also provide protection from diverse biotic and abiotic stressors. In the medfly, *Ceratitis capitata*, the primary components of the accessory gland secretion deposited during oviposition are ceratotoxins, which act as potent antibacterial agents (Marchini, Bernini, Marri, Giordano, & Dallai, 1991). Similarly, accessory gland secretions from the sand fly, *Phlebotomus papatasi*, have antimicrobial effects that protect the eggs, developing embryos, and adult female reproductive tract from microbial invasion (Rosetto et al., 2003). Some male seminal fluids also contain antimicrobial peptides, probably for similar reasons as in the female (Lung, Kuo, & Wolfner, 2001; Poiani, 2006). Female accessory gland proteins can also be a source of nutrition for developing progeny, either while growth occurs in the mother or as a secreted food source to nourish external progeny (Benoit, Kölliker, & Attardo, 2019; Benoit et al., 2015; Kaiwa et al., 2014; Kaulenas, 2012).

In this study, we use RNA-seq, proteomics, and functional analyses to examine the reproductive physiology of *B. antarctica*. Specifically, male and female accessory glands are examined to identify factors related to male accessory protein generation and synthesis of egg components during oviposition. Proteomic analysis of the gel secretion is used to identify its components, while comparative genomic analyses are used to identify orthologs of specific reproduction-associated genes in mosquitoes and midges. Functional studies reveal that stress impinging on late instar larvae impacts synthesis of gene products associated with reproduction, lowering both male and female reproductive success. Furthermore, we determine that the gel likely acts to prevent egg dehydration and serves as thermal buffer, preventing overheating of the eggs. Population growth modeling reveals that impaired fecundity from larval stress may reduce reproduction below population replacement levels. Our analysis shows that reproduction in the Antarctic midge is directly impacted by larval stress, and identifies novel roles for products manufactured by the female accessory gland. These studies confirm that an understanding of reproductive biology is critical for establishing how these Antarctic extremophiles are able to survive and proliferate in the challenging polar environment.

## 2 MATERIALS AND METHODS

### 2.1 Midge collections

Antarctic midges were collected from islands near Palmer Station (64°46′S, 64°04′W) in January 2007 and January 2017. Males and females were separated based upon the major morphological characters described previously (Convey & Block, 1996; Sugg et al., 1983; Usher & Edwards, 1984), homogenized on-site, and stored in Trizol (Invitrogen) at -70 °C for shipment to the University of Cincinnati. Female and male accessory glands were also dissected (N = 20-30) and stored in Trizol similar to whole body stages.

Larvae were collected from the same location as adults. Larvae within organic debris were returned to Palmer Station, and larvae were extracted into ice water with a modified Berlese funnel. Following recovery, larvae were stored with substrate from their natural habitat (rocks, soils, moss, and the alga *Prasiola crispa*, which serves as a food source for *B. antarctica*) at 2-4 °C. Larvae were shipped to the University of Cincinnati and stored under similar conditions until they were used in studies examining the impact of larval stress or gel presence on adult fertility or egg viability, respectively.

### 2.2 RNA extraction and processing

RNA was extracted from the midges by homogenization (BeadBlaster 24, Benchmark Scientific) in Trizol reagent (Invitrogen), using manufacturer’s protocols with slight modification based on other studies of invertebrates (Hagan et al., 2018; Rosendale, Dunlevy, McCue, & Benoit, 2019). Extracted RNA was treated with DNase I (Thermo Scientific) and cleaned with a GeneJet RNA Cleanup and Concentration Micro Kit (Thermo Scientific) according to manufacturer’s protocols. RNA concentration and quality were examined with a NanoDrop 2000 (Thermo Scientific).

Poly(A) libraries were prepared by the DNA Sequencing and Genotyping Core at the Cincinnati Children’s Hospital Medical Center. RNA was quantified using a Qubit 3.0 Fluorometer (Life Technologies). Total RNA (150-300 ng) was poly(A) selected and reverse transcribed using a TruSeq Stranded mRNA Library Preparation Kit (Illumina). An 8-base molecular barcode was added to allow for multiplexing and, following 15 cycles of PCR amplification, each library was sequenced on a HiSeq 2500 sequencing system (Illumina) in Rapid Mode. For each sample, 30-40 million paired-end reads at 75 bases in length were generated. Raw RNA-seq data have been deposited at the National Center for Biotechnology Information (NCBI) Sequence Read Archive: Bio-project PRJNA576639. Along with the RNA-seq samples collected for this study, larval (control, dehydration, and cryoprotective dehydration) samples were acquired from Teets et al.(Teets et al., 2012) under the NCBI Bioproject PRJNA174315.

RNA-seq reads were trimmed for quality (Phred score limit of 0.05) and sequences with ambiguities were removed. In addition, five and eight nucleotides were removed from the 5’ and 3’ ends, respectively, and sequences shorter than 45 bases were removed. Reads before and after cleaning and trimming were examined with FastQC for quality (S. Andrews http://www.bioinformatics.babraham.ac.uk/projects/fastqc) to verify the quality of each set.

### 2.3 Gene expression analyses

RNA-seq analyses were conducted using two distinct pipelines. The first method utilized was CLC Genomics (Qiagen), as previously described (Hagan et al., 2018; Rosendale et al., 2019). Briefly, reads were mapped to contigs with a cutoff of at least 80% of the read matching at 90% identity with a mismatch cost of 2. Each read was permitted to align to only 20 contigs. Expression values were based on total read counts in each sample calculated as transcripts per million reads mapped. EdgeR was used to test significance among samples. A multiple comparison correction was performed (false discovery rate, FDR). Genes were considered to be differentially expressed if the fold change was greater than 2.0 and the p-value was < 0.05. In addition, genes were required to have at least 5 mapped reads per sample to be retained for further analyses. For whole-carcass expression analyses, genes were considered sample-specific if enrichment was noted in relation to both of the other whole-carcass datasets. For accessory gland specific analyses, genes were considered tissue specific if they were enriched relative to the relevant whole-carcass dataset (female or male). In addition, a *de novo* assembly was conducted on the female accessory gland RNA-seq datasets using Trinity (Grabherr et al., 2011) based on standard methods to determine if bacterial symbionts were present in this organ and could be transferred to the egg while in the gel.

The second method for examining transcript expression involved utilization of RNA-seq tools available through the Galaxy software package (www.https://usegalaxy.org/) (Afgan et al., 2018; Goecks, Nekrutenko, & Taylor, 2010) using Salmon with the suggested settings. Differential expression between genes was examined with the DeSeq2 package (Love, Huber, & Anders, 2014). A general linearized model assuming a binomial distribution followed by a false discovery rate (FDR) approach were utilized to account for multiple testing. Cut-off values for significance, enrichment and sample-specificity were the same as those used in analysis conducted with CLC Genomics.

Transcripts identified as sex- or development-specific were examined using the CLC-based pipeline as there was over 95% overlap between each RNA-seq analysis method. Pathways enriched within males, females, and larvae were identified with a combination of Database for Annotation, Visualization and Integrated Discovery (DAVID (Huang da et al., 2009)), Blast2GO enrichment analyses(Conesa et al., 2005), CLC gene set enrichment analysis (Clark & Ma’ayan, 2011), and g:Profiler (Raudvere et al., 2019). Due to the taxonomic limitations of DAVID, sets of enriched transcripts were compared by BLASTx to the *An. gambiae* and to the *D*. *melanogaster* RefSeq protein datasets to identify homologous sequences. Blast hits (e-value < 0.001) from these two species were submitted to DAVID. There was considerable overlap between the results, and only the CLC-based methods were used in subsequent analyses.

Lastly, we utilized weighted correlation network analysis (WGCNA) to identify specific modules of genes that have similar expression profiles across developmental stages and accessory glands (Langfelder & Horvath, 2008). WGCNA was implemented in an R software package used to construct correlation networks and describe correlation between gene expression across samples in RNA-seq or microarray data. Genes sharing similar patterns of expression across samples were clustered into modules to identify groups of biologically significant genes that were particular to one of the sample groups. For this analysis, RNA-seq data were screened for genes of zero variance prior to WGCNA, leaving 13,424 genes for signed network construction. The minimum module size allowed was 20 and the soft power was set to 14 as determined by the package’s scale-free topology function. Modules exhibiting the highest Pearson correlation coefficient were selected for further analysis to determine function and relationship to developmental stages and accessory glands. Modules identified as enriched in a specific developmental stage or tissue were examined for enriched GO categories with the use of g:Prolifer and DAVID, as described earlier.

### 2.4 Comparative analyses with other chironomid midges and *Anopheles* mosquitoes

This is the first study to examine genome-wide, sex- and stage-specific expression in a midge. However, a recent study examines sex-specific expression in four species of anopheline mosquitoes, a clade not distantly related to midges, along with expression specific to the male and female reproductive tracts (MRT and FRT, respectively (Papa et al., 2017)). The species covered in this study were *An. gambiae, An. minimus, An. albimanus,* and *An. arabiensis.* In addition, male accessory glands enriched genes for *B. antarctica* were directly compared to those from Anopheles male accessory glands(Izquierdo et al., 2019). Predicted gene sets from this study that had enriched expression in specific stages and organs were used in comparative analyses with *B. antarctica.* The genomes of four species of chironomid midges were also acquired: *Clunio marinus* (Kaiser et al., 2016)*, Parochlus steinenii* (Kim et al., 2017)*, Polypedilum vanderplanki* (Gusev et al., 2014), and *Polypedilum nubifer*(Gusev et al., 2014). Studies that resulted in these sequenced midge genomes did not include analyses of differential expression between sexes.

For comparative analyses between midges, predicted gene sets for *B. antarctica* were compared with genomes of each midge species, and results of these four analyses were pooled to establish putative sets of genes common to all five species. This analysis also resulted in identification of differentially expressed genes unique to *B. antarctica*. Similarly, predicted gene sets from *B. antarctica* were compared with each species of *Anopheles* mosquito. Antarctic midge gene sets were compared to *Anopheles* whole carcass gene sets, as well as reproductive tract-specific gene sets (Papa et al., 2017). Lastly, we compared the expression of genes within the male accessory glands to orthologous gene sets that are uniformly expressed in the male accessory glands of mosquitoes (Izquierdo et al., 2019). The common gene sets produced by these analyses were then subjected to ontological analyses, using gProfiler, to establish sex-specific enriched pathways. tBLASTp analyses (e-value < 0.001) were performed using CLC Genomics Workbench (CLC bio Qiagen). Protein sequences were defined as orthologs if they were reciprocal-best BLASTp hits having an e-value < 10^−10^. Overlap was compared between these analyses to produce putative sex-specific transcript sets.

Transcription factors (TFs) and their predicted DNA binding motifs were identified based on methods used for other invertebrate genomes (Attardo et al., 2019; Benoit et al., 2016; Olafson et al., 2019). In brief, putative TFs were identified by scanning the amino acid sequences of all proteins for putative DNA binding domains using the HMMER software package (Eddy, 2009) and a compilation of Pfam DNA binding domain models (Weirauch & Hughes, 2011). Experimentally determined DNA binding motifs were then inferred from other species (*e.g., Drosophila*) based on amino acid identity, using previously established rules (Weirauch et al., 2014). Using this collection of inferred DNA binding motifs, we examined enrichment of each motif within the 500 and 2000 bp promoter regions of genes with increased expression in each sex, larvae, and accessory glands. These results were then compared to the expression profiles of each TF to determine specific TF candidates that might regulate sex and reproduction associated genes.

### 2.5 PCR and qPCR analyses

Select genes of interest that were highly enriched within males and females were examined by PCR. Total RNA was extracted from males, females, larvae, female accessory glands, and male accessory glands as described previously in the RNA-seq section and used as a template for Superscript III reverse transcriptase according to the manufacturer’s protocols (Invitrogen). PCR was performed with gene-specific primer pairs (Table S15) using a DNA polymerase kit (Promega). The PCR conditions were 95 °C for 3 min, 35 cycles of 30 sec at 95 °C, 52-56 °C for 1 min, and 1 min at 70 °C using an Eppendorf Mastercycler Pro Series. Three independent (biological) replicates were conducted for each sex or tissue stage.

qPCR analyses were conducted based on previously developed methods (Rosendale, Romick-Rosendale, Watanabe, Dunlevy, & Benoit, 2016). RNA was extracted as described previously for independent biological replicates. Complementary DNA (cDNA) was generated with a DyNAmo cDNA Synthesis Kit (Thermo Scientific). Each reaction used 250 ng RNA, 50 ng oligo (dT) primers, reaction buffer containing dNTPs and 5 mmol•l^−1^ MgCl_2_, and M-MuLV RNase H+ reverse transcriptase. KiCqStart SYBR Green qPCR ReadyMix (Sigma Aldrich, St Louis, MO, USA) along with 300 nmol l^−1^ forward and reverse primers, cDNA diluted 1:20, and nuclease-free water were used for all reactions. Primers were designed using Primer3 based on contigs obtained from the transcriptome analysis (Table S15). qPCR reactions were conducted using an Illumina Eco quantitative PCR system. Reactions were run according to previous studies(A. J. Rosendale et al., 2016). Four biological replicates were examined for each sex, and three biological replicates were examined for each accessory gland. Expression levels were normalized to *rpl19* using the ΔΔCq method as previously described(Joshua B Benoit et al., 2018; A. J. Rosendale et al., 2016). Fold change was compared between larvae, males, females, and accessory glands followed by a Pearson correlation coefficient (r).

### 2.6 Proteomics analysis of accessory gland-derived gel

Samples were analyzed at the Proteomics and Metabolomics Laboratory at the University of Cincinnati. Two proteomic samples were collected by removing eggs from the gel with a micropipette and dissolving the gel in 1X PBS with 0.1% Tween. Samples (4 µg) were run on a 1D SDS PAGE gel and silver stained to confirm the presence of proteins; at least 15 distinct proteins could be visualized. Based on this initial characterization, gel proteins (6 µg) were run 2 cm into a 1D 4-12% Bis-Tris Invitrogen NuPage gel using MOPS buffer. Lanes were excised, reduced with 10 mM dithiothreitol, alkylated with Iodoacetamide and digested with trypsin according to the standard protocol (Heaven et al., 2016; Turnier et al., 2018). The resulting peptides were concentrated with a speed vac centrifuge and resuspended in 0.1% formic acid. Each sample (2 µg) was used in subsequent analyses. Nanoscale LC-electrospray ionization-MS/MS (nanoLC-ESI-MS/MS) analyses were performed on a TripleTOF 5600 (Sciex, Toronto, ON, Canada) coupled to an Eksigent (Dublin, CA) nanoLC ultra nanoflow system. Protein from each gel sample was loaded and analyzed as described (Heaven et al., 2016; Turnier et al., 2018).

The data were recorded using Analyst-TF (v.1.6) software and searched against the *B. antarctica* genome (Kelley et al., 2014) using the Protein Pilot program (Sciex). Gel proteins were compared to those with differential expression in specific tissues from our RNA-seq studies. Protein, carbohydrate, and lipid content were examined through spectrophotometric assays based upon methods described in Rosendale et al. (2019)

### 2.7 Thermal buffering by the accessory gland gel

The gel has been suggested to serve as a source of nutrients for newly emerged larvae (Convey, 1992; Convey & Block, 1996; Harada et al., 2014). In addition to this role, we examined whether the gel increases thermal buffering capabilities of the egg compared to eggs directly deposited on the local substrate. To examine the effect, we placed an Omega thermocouple within six gels and immediately adjacent to these six gels at a field location near Palmer Station. Temperature was measured every minute over the course of three days.

To establish whether the gel prevents egg death caused by thermal stress, eggs with and without gels were exposed to 20 °C for two hours. Females were allowed to lay their eggs onto filter paper disks (Whatman) which were placed in 50 ml centrifuge tubes before being transferred to a 4 °C water bath. Temperature was then ramped up to 20 °C over the course of four hours 4 °C per hour before slowly being reduced back to 4 °C over the same course of time. These samples were compared to those that were held statically at 4 °C without the gel. Following treatment, all eggs were maintained at 4 °C and monitored for larval emergence.

### 2.8 Role of the accessory gland gel in dehydration

To determine if the accessory gland gel could prevent egg dehydration, we subjected midge eggs to dehydrating conditions with and without the gel’s presence. All eggs were removed from the gel with a fine metal probe and half were carefully reinserted into the gel. Three groups of eggs with or without the gel were moved to 75% RH at 4 °C for 12 hours. Following this treatment, eggs with no gel were placed back into the gel. Viability was determined by counting the number of larvae that emerged from the total number of eggs.

### 2.9 Impact of larval dehydration stress on fecundity

To determine if larval dehydration stress impacts adult fecundity, we performed combined RNA-seq analysis (described previously) and exposure of larvae to dehydration stress, both standard and cryoportective (Teets et al., 2012). RNA-seq studies were conducted on larvae that were quickly dehydrated (30% water loss) and on those that had undergone a slower form of dehydration, cryoprotective dehydration (30% water loss). The resulting data sets were then examined to find genes with increased expression in male or female accessory glands.

To determine whether mating is directly impacted by dehydration stress, groups of 100 fourth instar larvae (final larval instar) were held at 75% RH until they lost 40% of their water content. Following dehydration, larvae were returned to the standard rearing conditions and monitored every 12 hours for the presence of pupae or newly emerged adults. Each adult was removed and stored separately at 98% RH, 4°C until mating, which occurred no later than four days after treatment. The following mating pairs were examined: males from dehydrated larvae vs. control females, females from dehydrated larvae vs. control males, and both males and females from dehydrated larvae. Males and females that failed to copulate were removed from the experiment.

### 2.10 Statistics

Replicates are independent biological samples. Sample sizes are listed in each method section or the figure legend. Significance is indicated within each figure and/or in the figure legend. Statistical tests are listed within the respective section in the methods or in the figure legends. All statistical analyses were performed using JMP version 11 (SAS) or R-based packages.

### 2.11 Population level effects

To explore population level effects of dehydration, gel surrounding the egg mass, and thermal stress we used a Leslie matrix approach (Lefkovitch, 1965). Here, the dominant eigenvalue of the matrix is the population growth rate (λ). We simplified life history to egg to larvae to adult despite there being four larval instars, potentially occurring over several years. For control populations we used a mean fecundity of 42.9 (eggs laid), an egg survival rate of 0.82, and larval survival rate of 0.78. To determine the effects of larval dehydration on population growth, we used fecundities (eggs laid) of 34.6, 35.6, and 21.5 for male, female, and male and female dehydration, respectively; egg and larval survivorship were assumed to be the same as control populations. To determine the effects of gel on population growth we used all parameters (as for control populations), but reduced larval survival to 0.5, based on the experimental data from our studies. Similarly, to investigate effects of thermal stress we reduced egg survivorship to 0.63, but left all other parameters at control values. We also investigated a worst case scenario with male and female dehydration, no gel, and thermal stress. All values were derived from the previously described experiments. We determined the dominant eigenvalue of each matrix using the function “eigen” in R.

## 3 RESULTS

### 3.1 Description of mating and reproductive organs in *B. antarctica*

Beyond copulation and sex ratios (Edwards, & Baust, 1981; Harada et al., 2014; Sugg et al., 1983), little is known about mating and reproductive aspects for *B. Antarctica.* In this study, the reproductive organs were observed by dissection. Following copulation (Fig. 1A), a spermatophore containing sperm and other, likely accessory gland products, are transferred as a bundle to the females (Fig. 1A, inset). Females deposit a gel around the fertilized eggs, and it is within this gel that embryogenesis occurs (Fig. 1B). The source of the gel appears to be the female’s accessory gland (Fig. 3C) because the gland is depleted followed egg laying (Fig. 1D,E). The male reproductive organs, including the testes and accessory glands, are depicted in Fig. 1F. The general organization and structure of the male and female reproductive anatomy is similar to that of another midge, *Chironomus plumosus* (Wensler & Rempel, 1962) and mosquitoes (Masci et al., 2015), with the exception that the female accessory gland of *B. antarctica* is dramatically enlarged. It is this enlarged gland that we consider to be the source of the gel deposited around the eggs.

**Figure 1:**
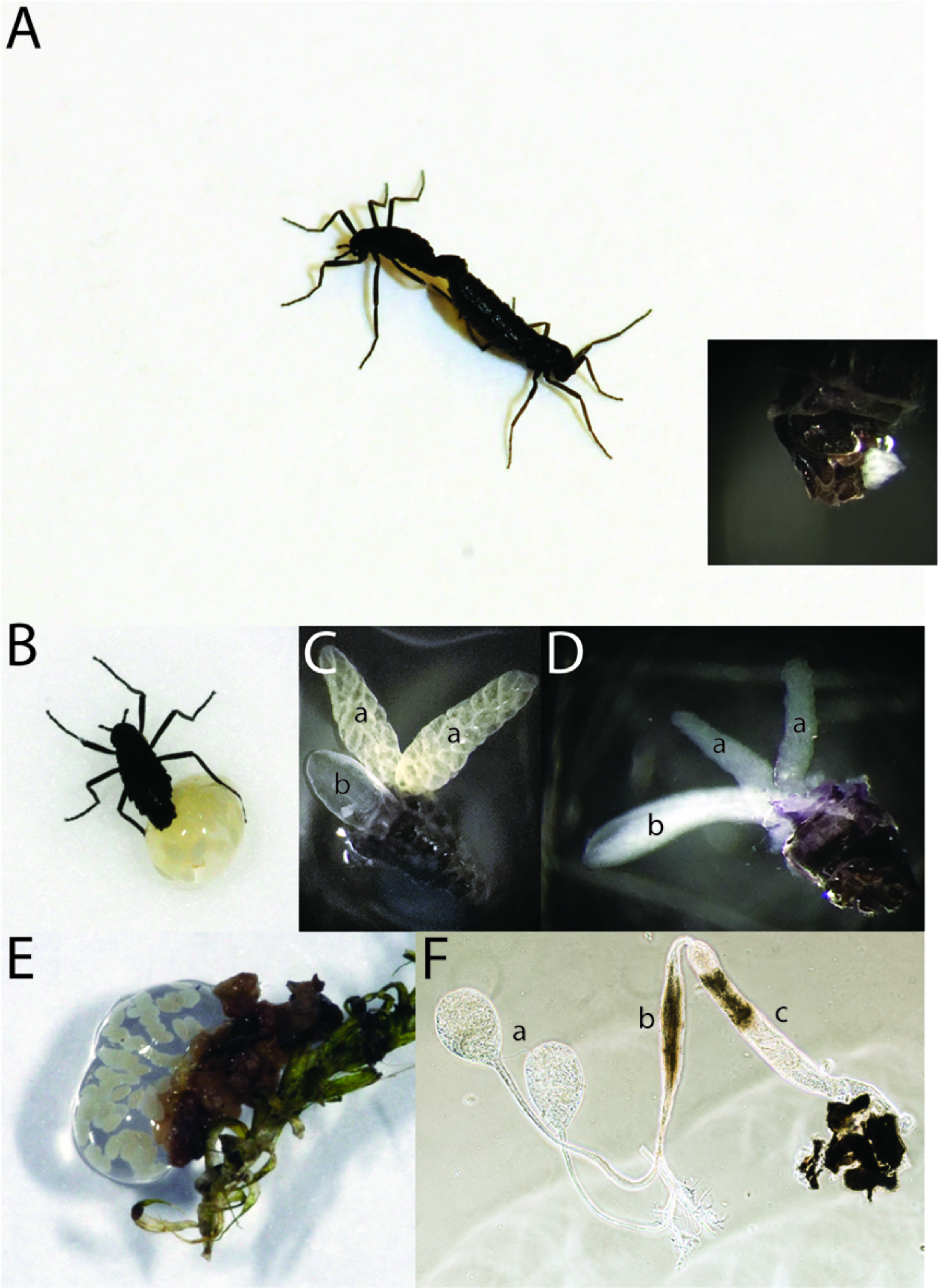
Antarctic midge, *Belgica antarctica*, during reproduction. A. Mating pair, male on left. Inset, spermatophore transferred to females immediately after copulation. Image is posterior end of female and white material is the spermatophore. B. Female depositing eggs and accessory gland-derived gel. C. Accessory gland (left middle, b) and ovaries (top left and right, a) of gravid females 3 days after adult eclosion. D. Female accessory glands (left, a) and ovaries (top and right, b) following egg and gel deposition. E. Egg mass following the completion of deposition. F. Male reproductive tract, a. testes, b, accessory gland, and c, common duct.

### 3.2 RNA-seq analyses of *Belgica* reproduction

Differential transcript levels were determined for males, females, and larvae using two independent pipelines using the *B. antarctica* genome as a reference (Kelley et al., 2014). The two pipelines yielded 95% overlap at our significance cut-off (two-fold difference, combined transcripts per million (TPM) among all samples of at least 5, and a correction-based P-value < 0.05). Between 60% and 71% of the reads from each RNA-seq set mapped to predicted genes. At the 4-fold or greater level for stage-specificity, overlap between the two methods was over 99%; thus we used the CLC Genomics (Qiagen) pipeline (Attardo et al., 2019; Rosendale et al., 2019) for the remaining analyses (Table S1). When expression differences were compared among all developmental stages, each sex and associated reproductive organs were most similar, followed by similarity between adults, while the larval stages were the most unique (Fig. 2). When the *de novo* assembly was examined for presence of putative microbial symbionts, we detected no substantial matches (Table S2), indicating the gel does not likely serve as a source of potential microbial symbionts as in other insects (Kaiwa et al., 2014).

**Figure 2:**
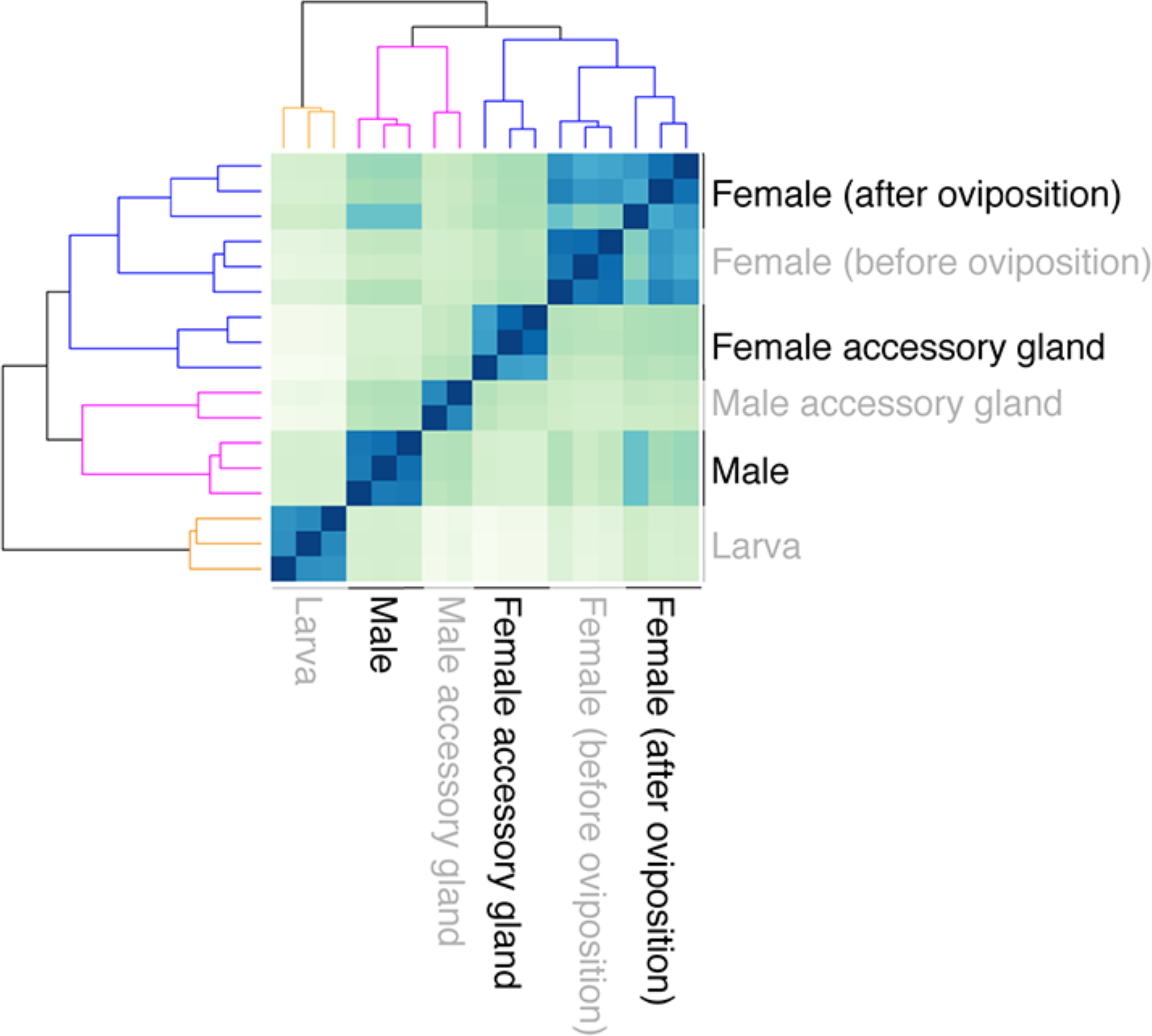
Gene expression heat map of Antarctic midge, *Belgica antarctica*, during development, between sexes, and for accessory glands. Hierarchical clustering of RNA-seq gene expression patterns for males, females, larvae, and accessory glands based on sample distance (Euclidean distance matrix) of differentially expressed contigs.

**Figure 3:**
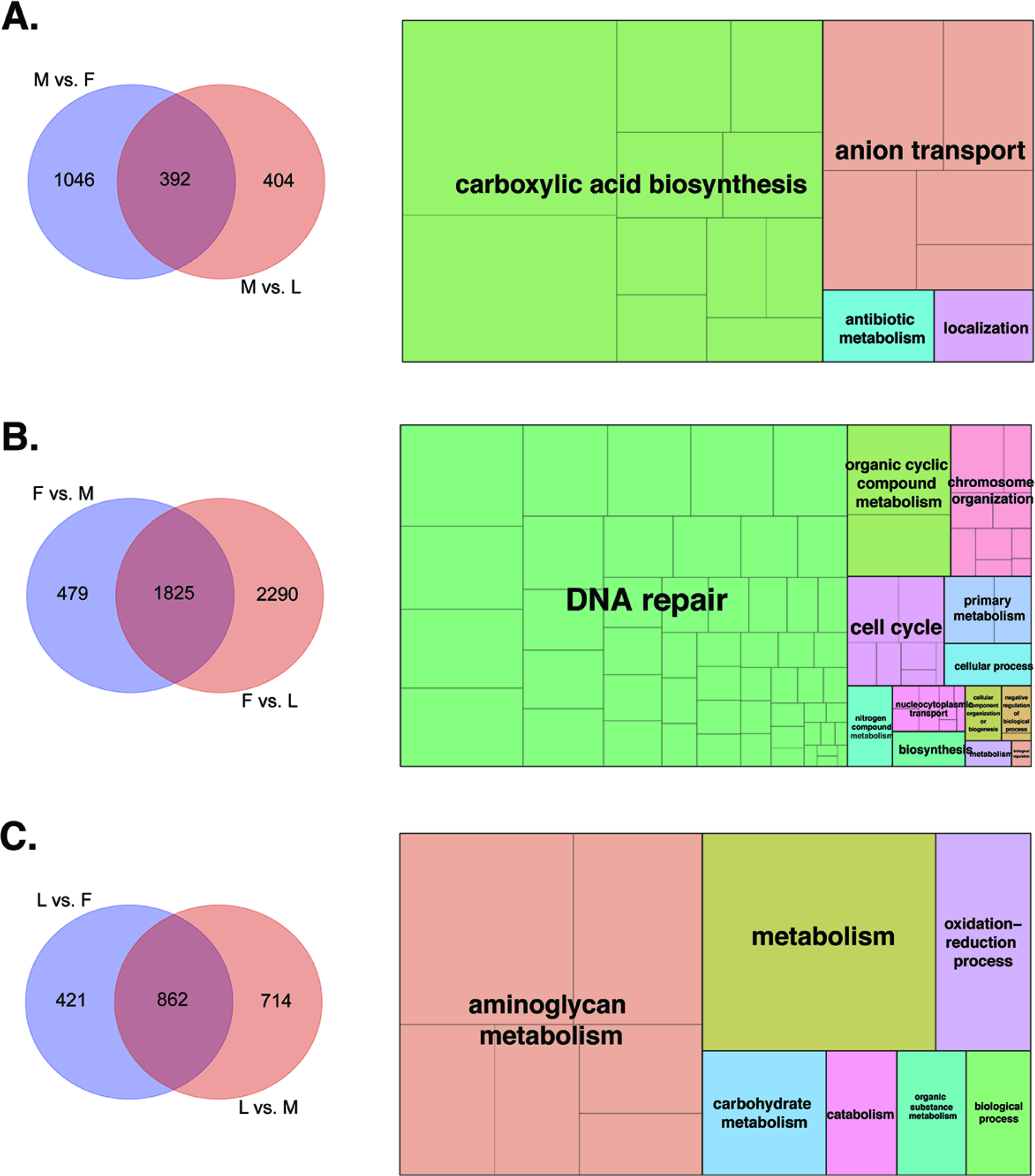
Genes uniquely enriched for the Antarctic midge, *Belgica antarctica* in males, females, and larvae and associated gene ontology enrichment. A. Gene enriched in males (left) and gene ontology (right), B. Gene enriched in females (left) and gene ontology (right), C. Gene enriched in larvae (left) and gene ontology (right). Each box represents a specific category and color represent major GO groups.

Individual comparisons between stages revealed 392 male-enriched genes, 1825 female-enriched genes, and 862 genes enriched in larvae (Fig. 2, Table S3-S5). Specific gene ontology (GO) categories were associated with each stage including carboxylic acid biosynthesis in males, DNA repair in females, and aminoglycan metabolism in larvae (Fig. 3; Table S3-S5). When accessory glands were specifically examined, 20 genes were enriched in the female accessory gland and had significantly higher expression in females compared to other stages (Fig 4A; Table S6). A similar analysis for the male accessory gland identified 25 enriched genes. GO categories associated with the female accessory gland were associated with glycosylation and mucin biosynthesis. Notably, similar GO categories were associated with gene products of the male accessory gland (Fig. 4B; Table S7).

**Figure 4:**
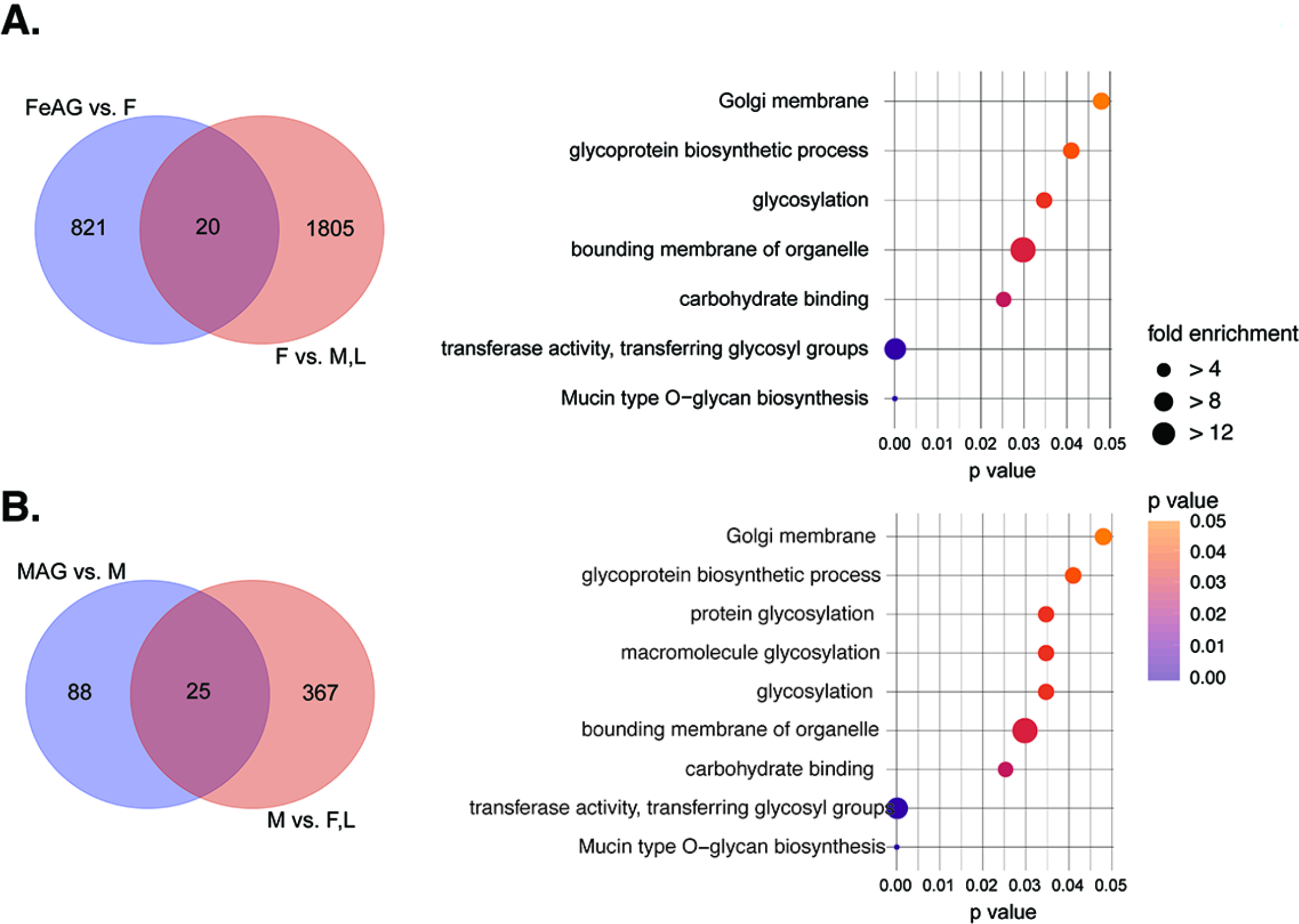
Genes uniquely enriched in the Antarctic midge, *Belgica antarctica*, female and male accessory glands and associated gene ontology enrichment. A. Genes enriched in female accessory glands (left) and gene ontology (right), B. Genes enriched in male accessory gland (left) and gene ontology (right). GO conducted with g:Prolifer (Raudvere et al., 2019).

Beyond this first analysis, weighted correlation network analysis (WGCNA, Langfelder & Horvath, 2008) was used to identify clusters (modules) of highly correlated genes between developmental stages and within specific reproductive organs (Fig. 5; Table S8-S12). Specific modules of enriched expression were identified in each tissue and developmental stage (Fig. 5B, C). When GO analyses were conducted for each developmental stage, unique GO categories were associated with larvae, females, and males (Fig. 5). Female accessory glands had few enriched GO categories based on WGCNA results; enriched categories included phosphatase binding, response to stimulus, and ion channel activity. Male accessory glands also had a low number of GO categories associated with enriched modules; these included metallopeptidase activity and integral components of the membrane. These results provide correlated stage- and tissue-specific expression modules. For larvae, males, and females, GO categories from stage specific modules showed overlap with our previous analysis (Fig. 3), but unique categories were also identified (Fig. 5).

**Figure 5:**
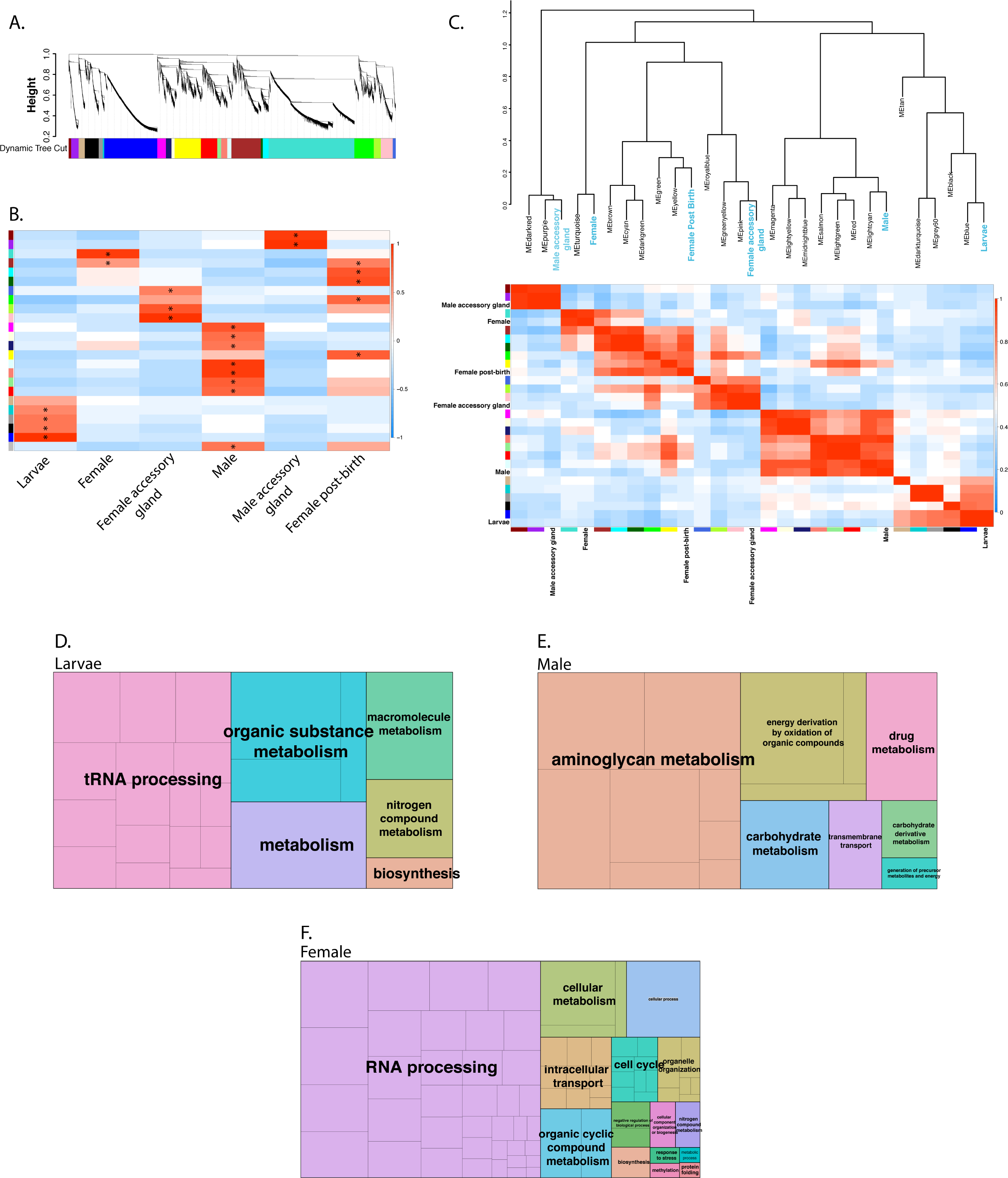
Weighted gene co-expression network analysis (WGCNA) across developmental stages and for male and female accessory glands of the Antarctic midge, *Belgica antarctica*. A. Average linkage hierarchical clustering dendrogram of the genes. Modules, designated by color code, are branches of the clustering tree. B. Correlation of module eigengenes to developmental and accessory gland traits. Each row corresponds to a module eigengene and columns are traits. *, represents values with a significant positive correlation for Pearson r (P < 0.05). C. Unsupervised hierarchical clustering heatmap (bottom) and dendrogram (top) of module eigengenes and traits. Gene ontology (GO) analysis of eigengenes associated with larvae (D), males (E), and females (F). GO conducted with g:Prolifer(Raudvere et al., 2019) and visualized with REVIGO(Supek, Bošnjak, Škunca, & Šmuc, 2011).

### 3.3 Proteomic analysis of accessory gland derived gel

Proteomic analysis of the gel surrounding the eggs revealed 24 associated proteins (Fig. 6). Three proteins comprised a majority of the gel (Fig. 6B; Table S13), including a vitellogenin-like protein (IU25_12621), larval serum protein (IU25_03947), and an apolipophorin (IU25_03809). The two analyzed gel samples had a highly-correlated protein content (Pearson correlation = 0.92, Fig. 3C). Expression analyses revealed that transcripts for most of the gel proteins are not directly generated in the female accessory gland (Fig. 3D), but instead are produced in other female organs (e.g. IU25_12621) or within the larvae (e.g. IU25_03947 and IU25_03809). qPCR validation of the protein gel components confirmed that transcripts for each of the gel proteins are expressed in tissues identified in the RNA-seq studies (Fig. 5E). Additional qPCR analyses for genes not involved in producing major components of the gel served as validation for results obtained from other RNA-seq samples (Fig. 5E).

**Figure 6:**
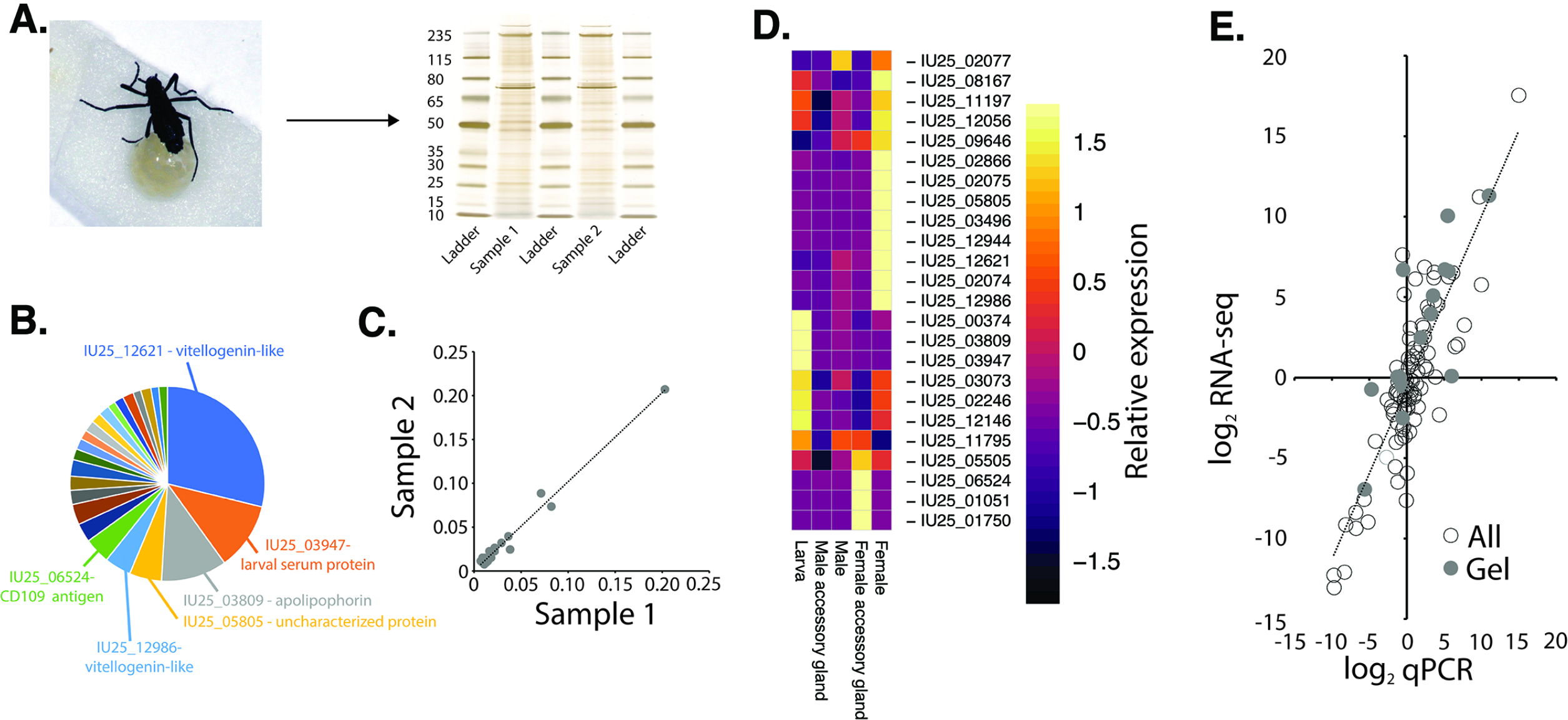
Proteomic analysis of female accessory gland derived gel material from the Antarctic midge, *Belgica antarctica*. A. Female depositing eggs with gel and protein components of two gel samples without eggs. B. Identification of proteins that represent at least 3% of total protein composition of the accessory gland gel. C. Congruence of protein abundance and content between two gel samples. D. Heatmap for transcript levels of gel-specific genes among larvae, females, males, and accessory glands. E. qPCR validation of RNA-seq data. All tested genes have Pearson correlation coefficients over 0.85. Gel specific genes have a Pearson correlation of 0.87.

### 3.4 Transcriptional regulation of reproductive-associated factors

To compare potential mechanisms regulating gene expression between samples, we predicted putative transcription factors (TFs, Fig. 7A, Table S14) and their DNA binding motifs. We next performed TF binding site motif enrichment analysis in the upstream regulatory regions of our differentially expressed gene sets (see Methods). This analysis revealed that fourteen TFs had significantly enriched binding sites in at least one gene set (Fig. 7B). Of these, one TF showed enhanced binding in the 500bp regulatory regions of male-enriched genes. The remaining thirteen showed enhanced binding in female-enriched genes; we limited our analysis to the set of seven TFs with enhanced binding in both 500 and 2000bp regions of the female- and/or FeAG-enriched gene sets. Five members of this set showed higher transcript levels in either the female or female accessory gland compared to other tissues and developmental stages (Fig. 7C). The increased gene expression in the respective tissues or stage along with enriched binding sites suggest that these TFS are key regulatory elements for female reproduction in *B. antarctica*. Transcription factors include forkhead box protein 1 and mothers against decapentaplegic (Mad) homolog 1. In *Drosophila* Mad has been identified as a participant in the signaling pathway of *decapentaplegic* (dpp), a morphogenetic protein that plays a role in regulating the development of egg polarity and early embryonic development (Xie & Spradling, 1998). Several transcription factors identified in our screening remain uncharacterized with no assigned biological functions.

**Figure 7:**
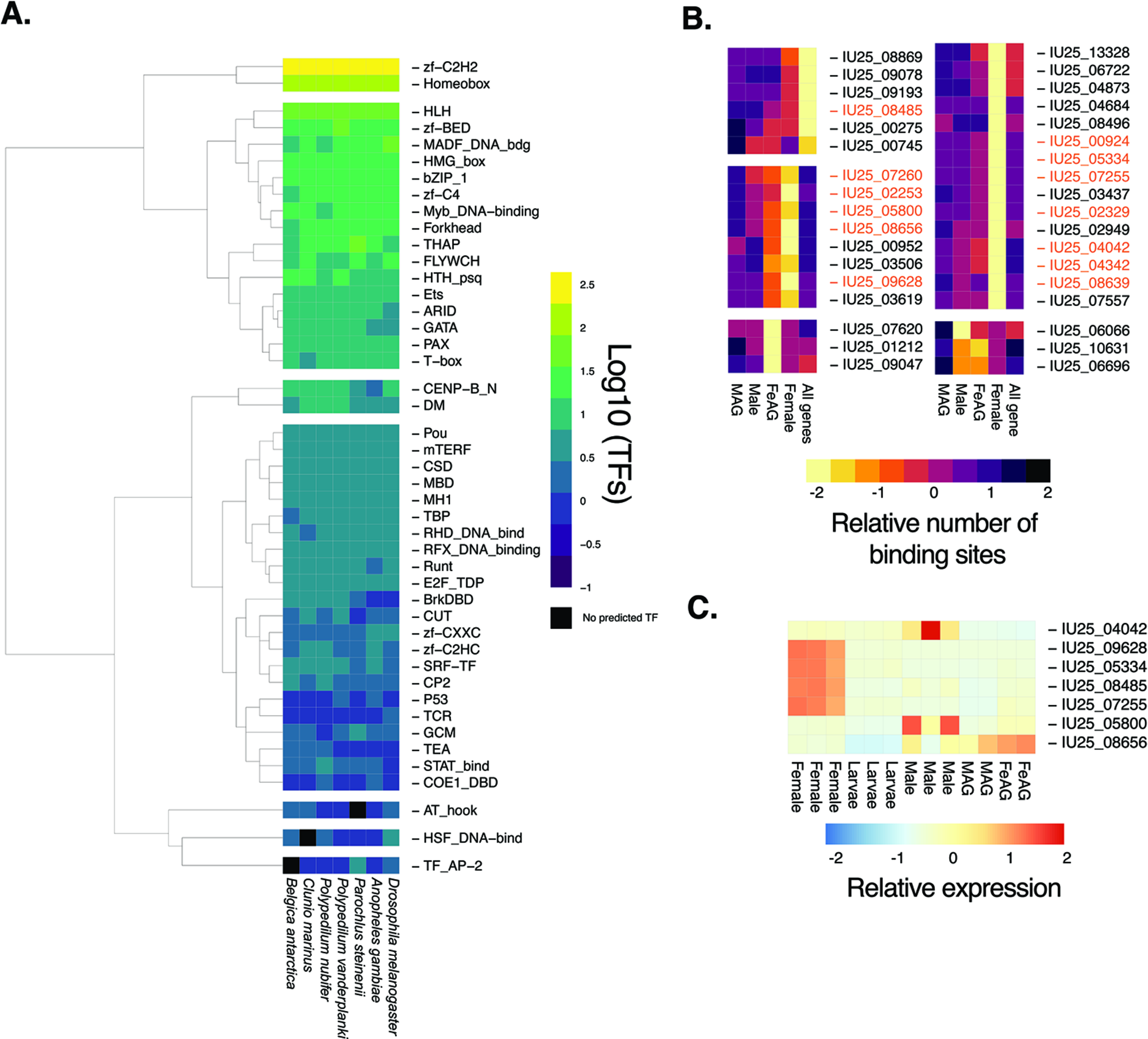
Transcription factors (TFs) and TF binding sites associated with reproduction in the Antarctic midge, *Belgica antarctica*. A. Relative abundance of transcription factors encoded by genomes of midges and mosquitoes. TF families towards the top contain more TFs in *B. antarctica*. B. Enrichment for specific TF binding site motifs in regulatory regions (2000 bp; 500 bp data not shown due to overlap with 2000 bp) of genes expressed highly in specific stages and accessory glands. Groups of TFs are separated by their motif enrichment profiles across samples. Those highlighted in orange are significantly enriched within the specific stage or tissue. Scale for heatmap is set at relative abundance on a Z scale of -2 to 2 across each row. C. Transcript levels of select TFs with significant motif enrichment in the promoters of genes expressed in specific tissues (orange font color in panel B).Scale for heatmap is set at relative abundance on a Z scale of -2 to 2 across each row. MAG, male accessory gland; FeAG, female accessory glands.

### 3.5 Comparative analyses between anopheline mosquitoes and chironomid midges

Sex- and tissue-specific gene sets from *B. antarctica* were compared with predicted gene sets from four species of *Anopheles* mosquitoes and four species of chironomid midges (Figs. 8 and 9). Additionally, *B. antarctica* enriched gene sets were compared with gene sets enriched in male and female reproductive tracts of the same four *Anopheles* mosquitoes (Papa et al., 2017) (Fig. 8 and 9) and in comparison to the male accessory gland enriched genes from five *Anopheles* species (Izquierdo et al., 2019)

**Figure 8:**
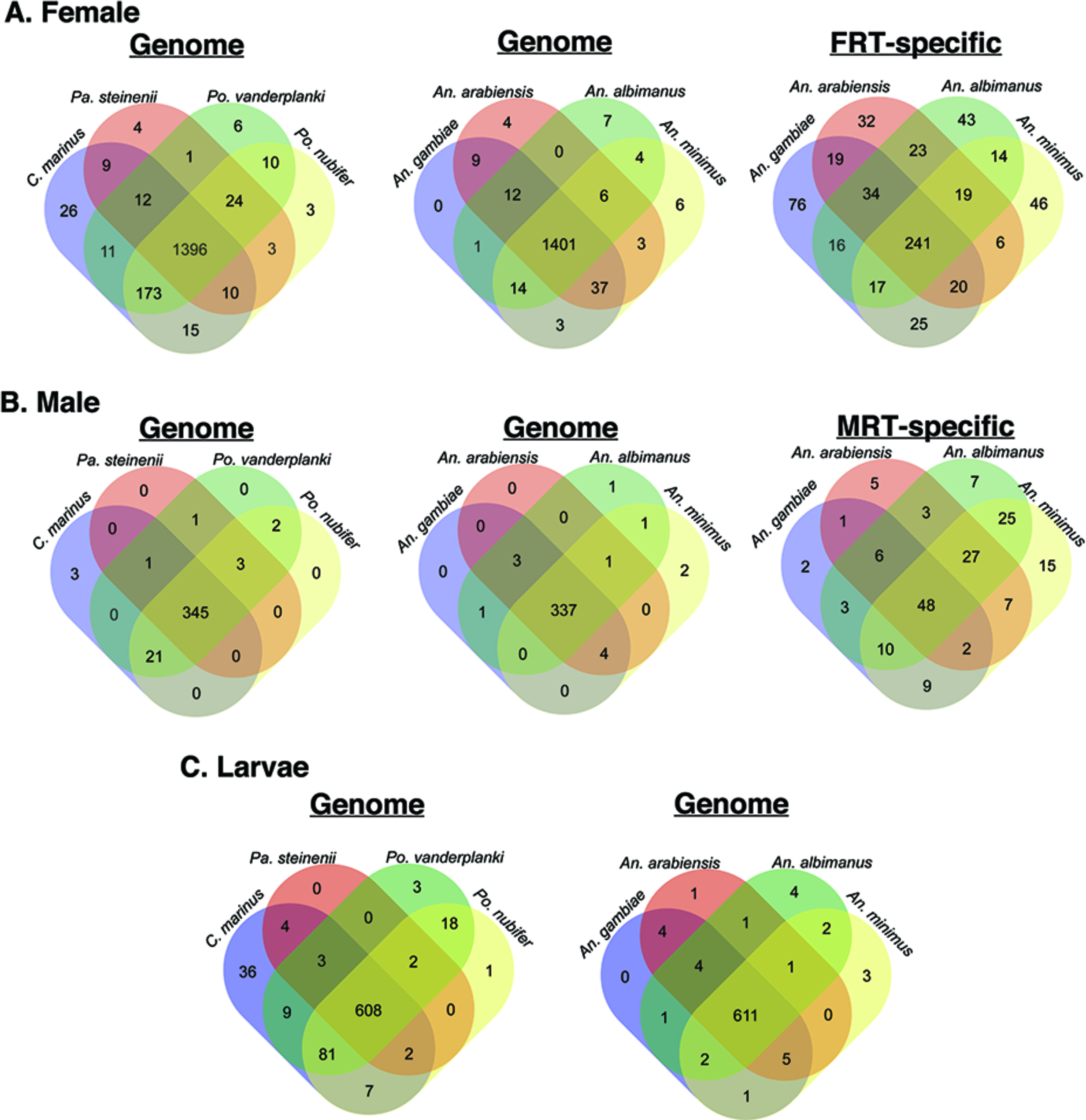
Comparative analysis of female, male, and larvae-specific gene sets with mosquitoes and midges to Antarctic midge, *Belgica antarctica*. A. Female-specific genes compared to midges (left), mosquitoes (middle), and genes with enriched expression in the female reproductive tract (FRT) of mosquitoes (right) (Papa et al., 2017). B. Male-specific genes compared to midges (left), mosquitoes (middle), and genes with enriched expression in the male reproductive tract (MRT) of mosquitoes (right)(Papa et al., 2017). C. Larvae-specific genes compared to midges (left) and mosquitoes (right). Protein sequences were defined as orthologs if they had reciprocal-best BLASTp hits with an e-value < 10^−10^.

**Figure 9:**
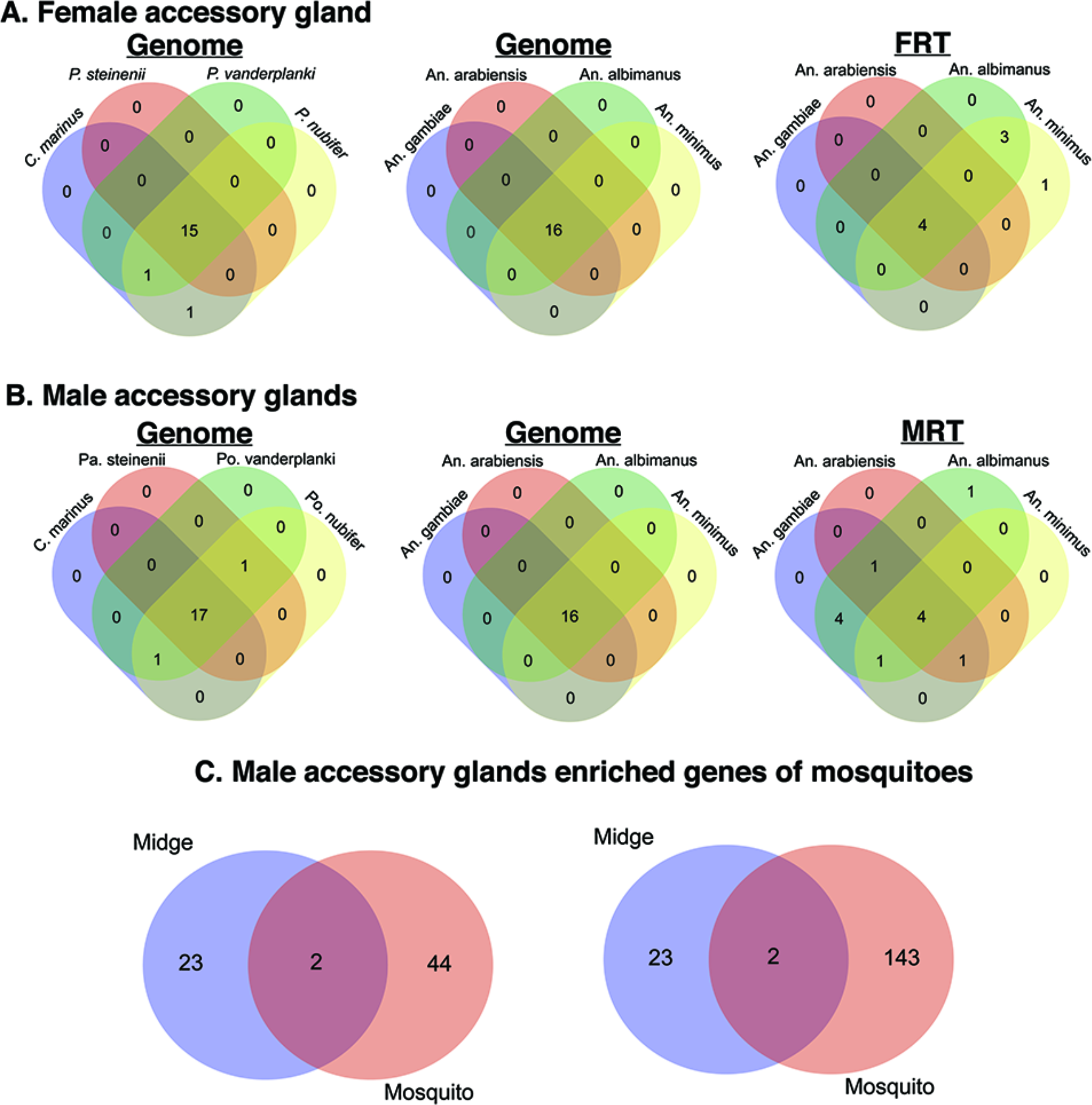
Comparative analysis of accessory gland gene sets with mosquitoes and midges to the Antarctic midge. A. Female accessory gland genes compared to midges (left), mosquitoes (middle), and genes with enriched expression in the female reproductive tract of mosquitoes (right) (Papa et al., 2017). B. Male accessory gland genes compared to midges (left), mosquitoes (middle), and genes with enriched expression in the male reproductive tract of mosquitoes (right) (Papa et al., 2017). C. Overlap between genes expressed in male accessory glands between mosquitoes and *B. antarctica*. Left, highly enriched in *Anopheles* male accessory gland. Right, enriched in *Anopheles* male accessory gland. Enrichment for *Anopheles* male accessory gland genes is based on values from Izquierdo et al. (2019). Protein sequences were defined as orthologs if they had reciprocal-best BLASTp hits with an e-value < 10^−10^.

Genes enriched in females of *B. antarctica* showed the highest degree of conservation among all species examined by a large margin. All five species of midges had 1396 genes in common, 1401 were shared with all four *Anopheles* species, and 1267 were common to all 9 species examined (Fig. 8). Gene ontology analysis of this set revealed enrichment in functions related to RNA, DNA, chromatin binding, nuclease activity, chromosome and organelle organization, transport of RNA and proteins in and out of the nucleus, RNA processing, and cell cycle regulation. No female-enriched genes detected in *B. antarctica* were unique when compared to the other eight species examined. *Belgica* females have 122 genes with no orthologues in any of the other four midge species. However, this gene set did not show significant enrichment in any GO functional categories.

Among males, 345 genes were common among midge species, and 337 genes were shared between *B. antarctica* and all four mosquito species (Fig. 8). Genes common to all midge species were enriched in functions associated with anion transport, alpha-amino acid catabolism, and carboxylic acid transmembrane transport. As with females, no genes were identified in the male-enriched gene set that were unique to *B. antarctica*. There were 16 genes unique to *B. antarctica* among the five midge species, but, again, they were not significantly enriched in any single functional category.

The larva-enriched gene set was compared to the midge and mosquito gene sets. Over 600 genes were identified with orthologues common to all examined midge and mosquito species (Fig. 8). The gene set common to all midge species was enriched in functional categories associated with chitin metabolism, iron ion binding, and cytochrome P450-driven metabolism. Two identified genes were unique to larvae of *B. antarctica* (Fig. 8; Table S5): one is uncharacterized, and the other putatively encodes polyubiquitin B. Gene ontology analysis comparing larvae of *Belgica* with other midges revealed 88 unique genes, but no functional categories were significantly enriched for this gene set.

Gene sets specifically enriched in male and female reproductive tracts of each mosquito species were compared to sex-specific gene sets in *B. antarctica* (Fig. 9; Table S6-7). Comparison of the female-enriched gene set with the female reproductive tract (FRT)-specific gene set yielded 241 genes with putative FRT orthologues. This set was particularly enriched in GO categories related to mitotic cell cycle processes and regulation of gene expression. For example, several orthologues in this set code for cyclins, cyclin dependent kinases, zinc-finger proteins, transcription factors, and transcription termination factors. Additionally, a large number of genes in this set (∼44%) mapped to apparently uncharacterized orthologs. The comparison between male-enriched and Male Reproductive Tract (MRT) specific genes yielded a common set of 48 genes. This set is dominated by CLIP-domain serine proteases and cytochrome p450s, and also contains three glucose-dehydrogenases (Fig. 9). When *B. antarctica* MAG-enriched genes are compared to those for mosquitoes, only four overlapping genes were identified based on moderate or high expression reported in a previous study(Izquierdo et al., 2019) of the mosquito MAG (Fig. 9C); these were identified as venom allergen, carboxypeptidase, serine protease, and a gene of unknown function.

### 3.6 Dehydration reduces larval serum protein and possibly subsequent egg production

Since specific female accessory gland components are expressed in larvae, we asked whether larval dehydration impacts expression of adult female accessory gland components (Fig. 10). To do so, we re-analyzed results from a previous RNA-seq study (Teets et al., 2012) on dehydration and cryoprotective dehydration of *B. antarctica* (Fig. 10). One of the major female accessory gland proteins (IU25_03947) had significantly reduced expression when larvae experienced either dehydration or cryoprotective dehydration (water loss specifically induced by cold temperatures) (Fig. 10A-B). This protein is a component of the accessory gel and, as a hexamerin, serves as a protein reserve (Burmester, 1999; Burmester, Massey Jr, Zakharkin, & Benes, 1998; Telfer & Kunkel, 1991). The dehydration-evoked suppression in transcripts associated with reproductive-associated factors possibly results in reduced amounts of materials invested in the progeny at birth and reduced egg output (Fig. 10D-F). These results suggest that stress experienced as larvae may have a direct impact on female reproduction, most likely acting through reduced expression of larval serum protein.

**Figure 10:**
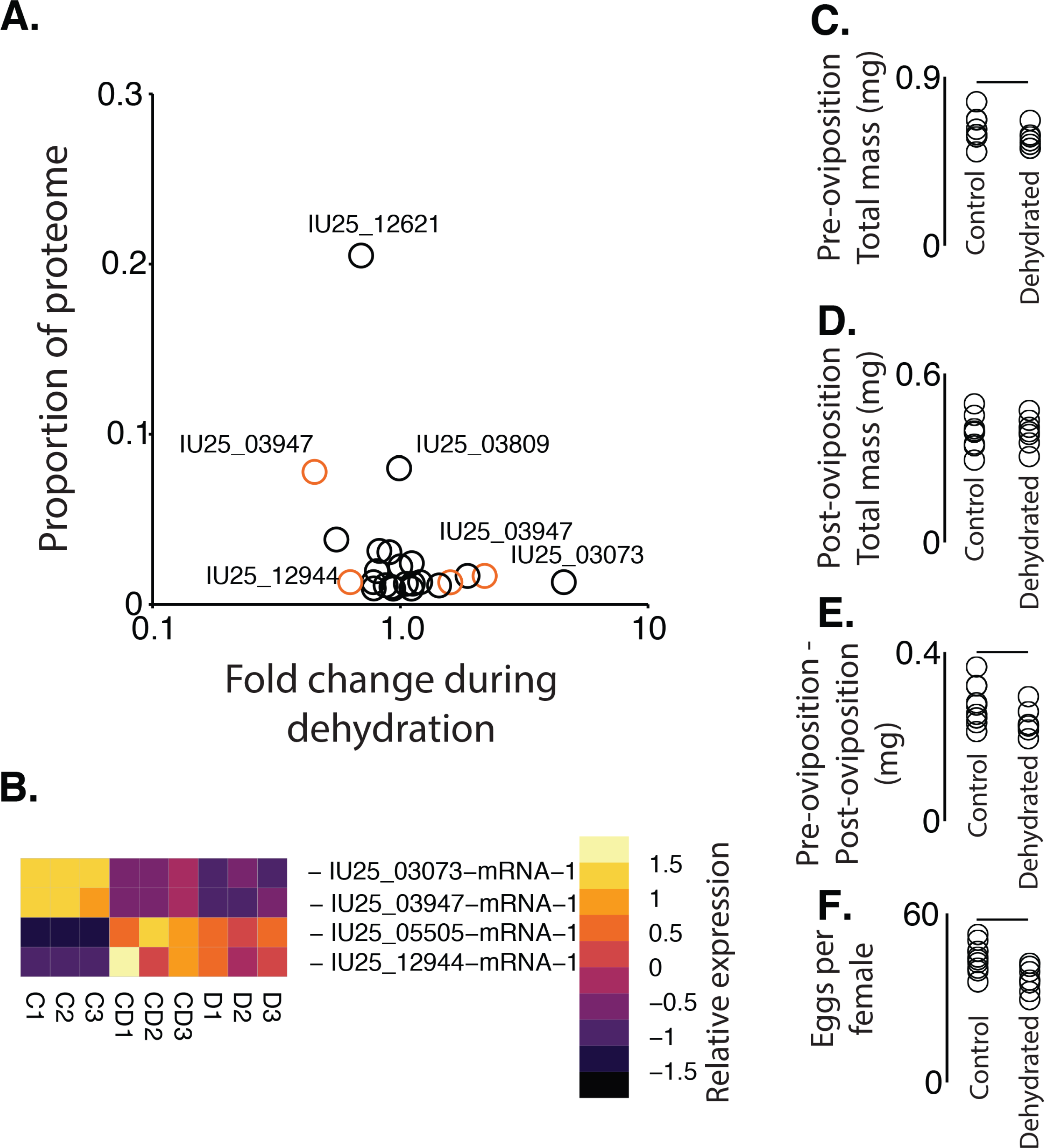
Expression changes in gel-associated proteins in larvae following dehydration stress. A. Transcript level changes in larvae for gel proteins following dehydration stress. RNA-seq studies were acquired from Teets et al. (2012). Orange denotes significance between control and dehydrated larvae based on RNA-seq analyses. B. Heat map of transcript levels for gel-associated proteins during dehydration (D) and cryoprotective dehydration (CD) compared to control (C) that are components of the gel proteome and significantly altered by dehydration. C. Total mass before eclosion, D. post-eclosion total mass, E. mass change after eclosion, and total egg production in females when control (non-dehydrated) and dehydrated larvae were allowed to complete development. Analysis of variance was utilized to examine statistical differences with the use of R statistic packages. Bars above indicate significance at P < 0.05.

### 3.7 Dehydration reduces specific male accessory gland components and impacts fertilization

We next examined the impact of larval dehydration stress on male fertility. To do so, we analyzed expression of accessory gland components in adult males after larval dehydration or cryoprotective dehydration (Fig. 11A). Thirteen genes enriched in male accessory glands were differentially expressed in the adult male following larval dehydration compared to fully-hydrated individuals (Fig. 11A). Enrichment of glutathione transferase activity and phosphatidylcholine metabolic processes were detected. Male body mass was not altered following dehydration stress, but mating was substantially compromised (Fig. 11B-C). When males dehydrated as larvae mated with unstressed females, fewer eggs were deposited, and additive reductions were noted if dehydrated males mated with females that were dehydrated as larvae (Fig. 11C). These results indicate an impact on male fertility brought about by dehydration stress during larval life.

**Figure 11:**
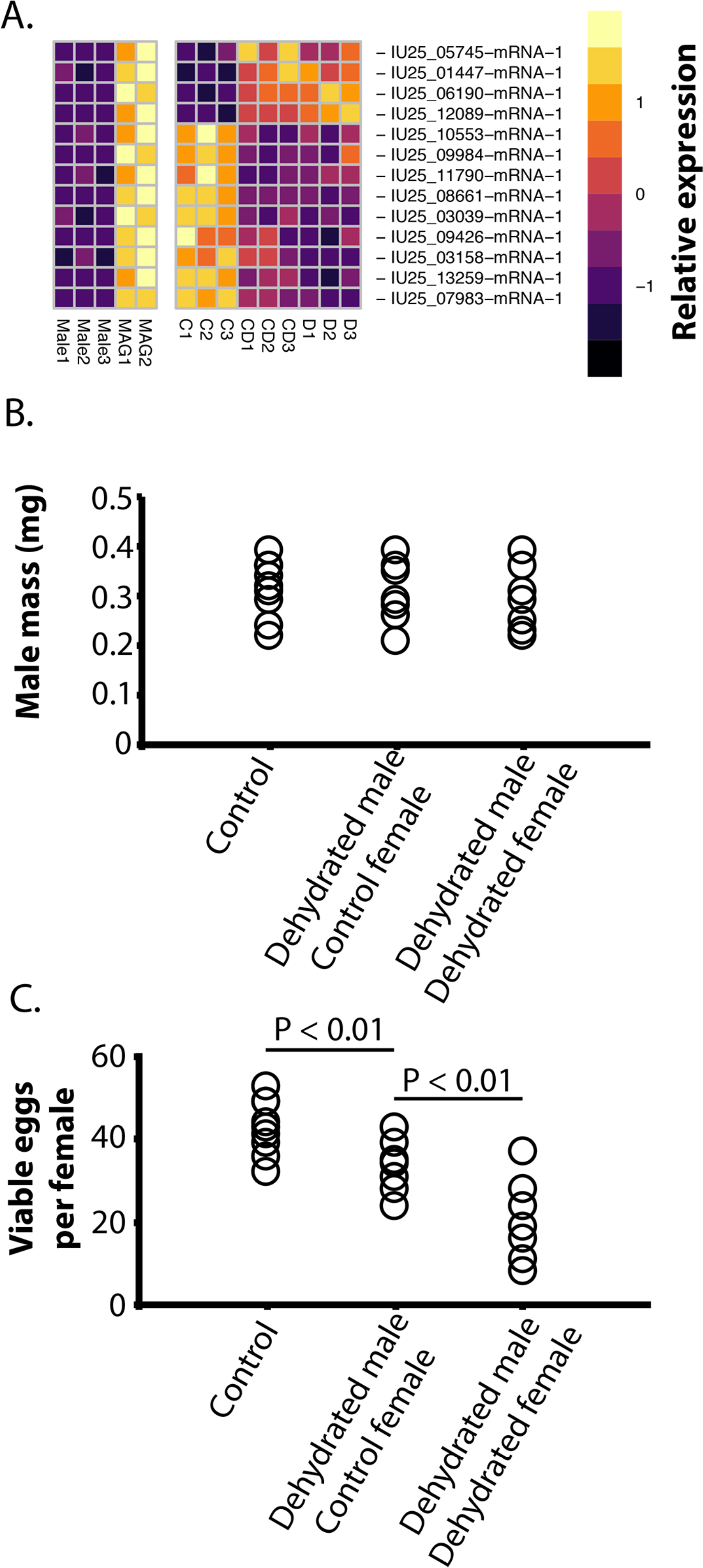
Impact of larval dehydration stress on male fertility. A. Expression profiles of male-associated genes with expression differences after larval dehydration (D, dehydration, C, control, and CD, cryoprotective dehydration, MAG, male accessory gland). B. Mass of males used in mating experiments from dehydrated or control larvae. C. Female fertility (control or dehydrated) following copulation with dehydrated or control males. Analysis of variance was utilized to examine statistical differences with the use of R statistic packages. Bars above indicate significance at P < 0.05 unless otherwise noted.

### 3.8 Role of gel as a nutrient source

Analyses of major gel protein components (over 3% of the total proteins of the gel) was performed. Amino acid content of the major protein components revealed that all essential amino acids can be provided within the gel (Fig. 12A). Predicted glycosylation and phosphorylation sites of the proteins suggest that these proteins could provide phosphate and sugar resources for the developing larva as the gel is consumed (Fig. 12). Spectrophotometric analysis of the gel revealed that 81%, 14%, and 5% of the caloric content consisted of proteins, carbohydrates, and lipids, respectively. Larvae were denied access to the gel to determine its impact on larval growth (Fig. 12B). Fewer of the gel-less larvae were alive after one month (Fig. 12B), and those remaining were nearly 30% smaller than larvae that hatched from eggs with free access to the gel (Fig. 12C). Importantly, these larvae were provided access to algae and other organic debris with and without gel, indicating that non-gel food resources do not fully compensate for loss of the gel as a food source. These results underscore the value of the gel for successful larval development.

**Figure 12:**
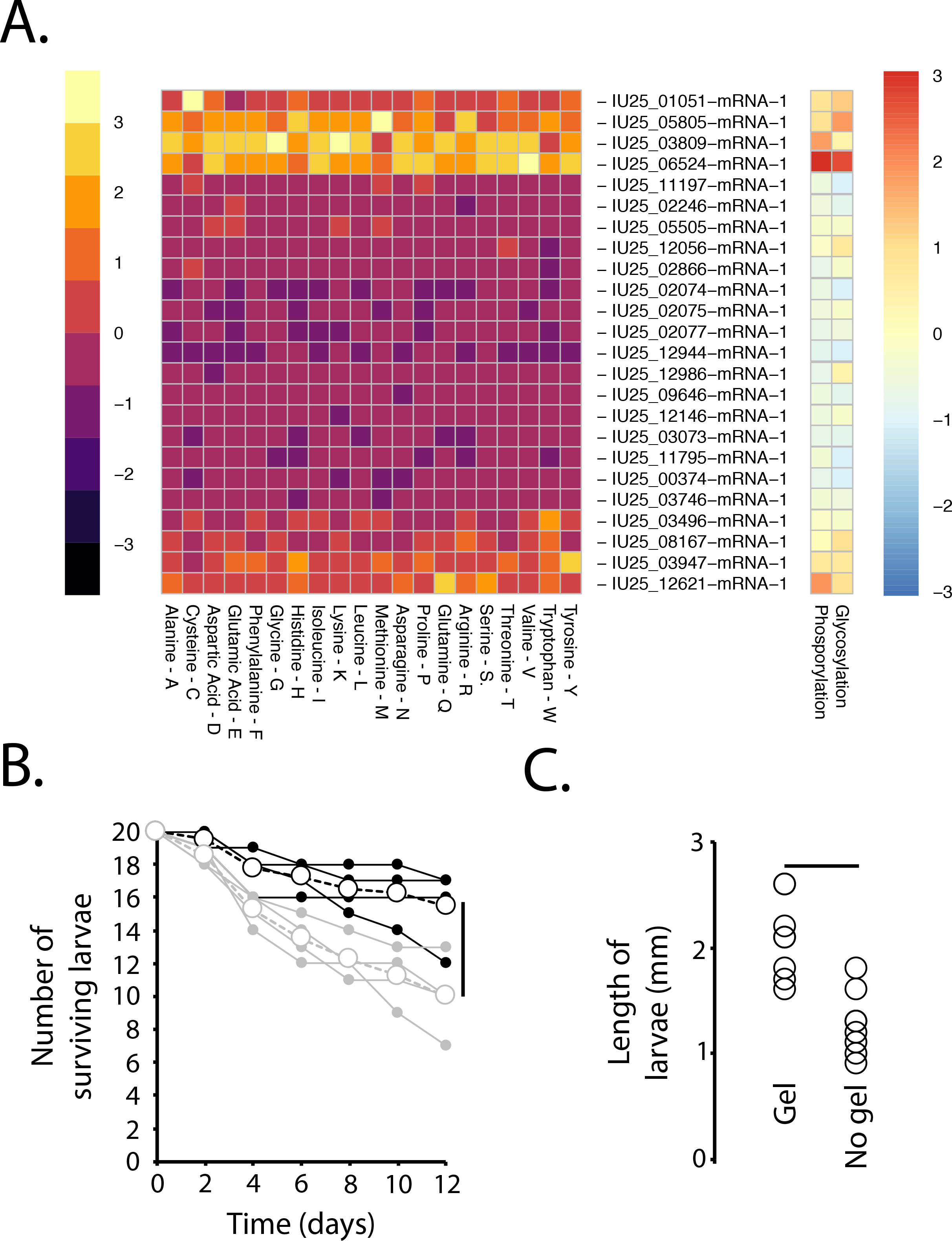
Accessory gland gel is critical for larval development. A. Amino acid composition and putative phosphate and glycosylation sites of gel proteins based on sequence information. Relative amounts are based on comparison levels between columns. B. Survival of developing larvae with (black) and without (gray) gel presence at larval ecdysis. Open circles are the average and filled circles are each replicate. C. Larvae length after 20 days with and without gel at larval ecdysis. Bar indicates significance at P < 0.05. Analysis of variance was utilized to examine statistical differences with the use of R statistic packages. Bars above or beside indicate significance at P < 0.05.

### 3.9 Thermal and dehydration buffering by the gel

We performed experiments to determine whether the gel impacts dehydration resistance in developing eggs. When gel-less eggs were held in desiccating conditions (75% relative humidity) at 4 °C for 12 h and subsequently treated with water, no viable eggs were detected, even when the eggs were subsequently covered with gel to promote growth (0% for all three replicates). By contrast, viability of eggs encased in gel exceeded 80% under in the same conditions, suggesting that the accessory gel protects eggs against dehydration-induced mortality. These results indicate that the gel is likely critical for maintaining water homeostasis within the egg and possibly for the developing embryo as well.

We also examined whether the gel acts as a thermal buffer. First, temperature changes were examined within the gel (next to the eggs) and on the surface immediately adjacent to the eggs (Fig. 13A). The gel buffered temperature changes during the course of the day by reducing both the maximum temperature and the rate of temperature change (Fig. 13). Gel-less eggs exposed to 20 °C displayed reduced viability when compared to eggs encased in gel and to those held constantly at 4 °C (Fig. 13). These results indicate that the accessory gland gel provides both thermal and dehydration protection to the eggs.

**Figure 13:**
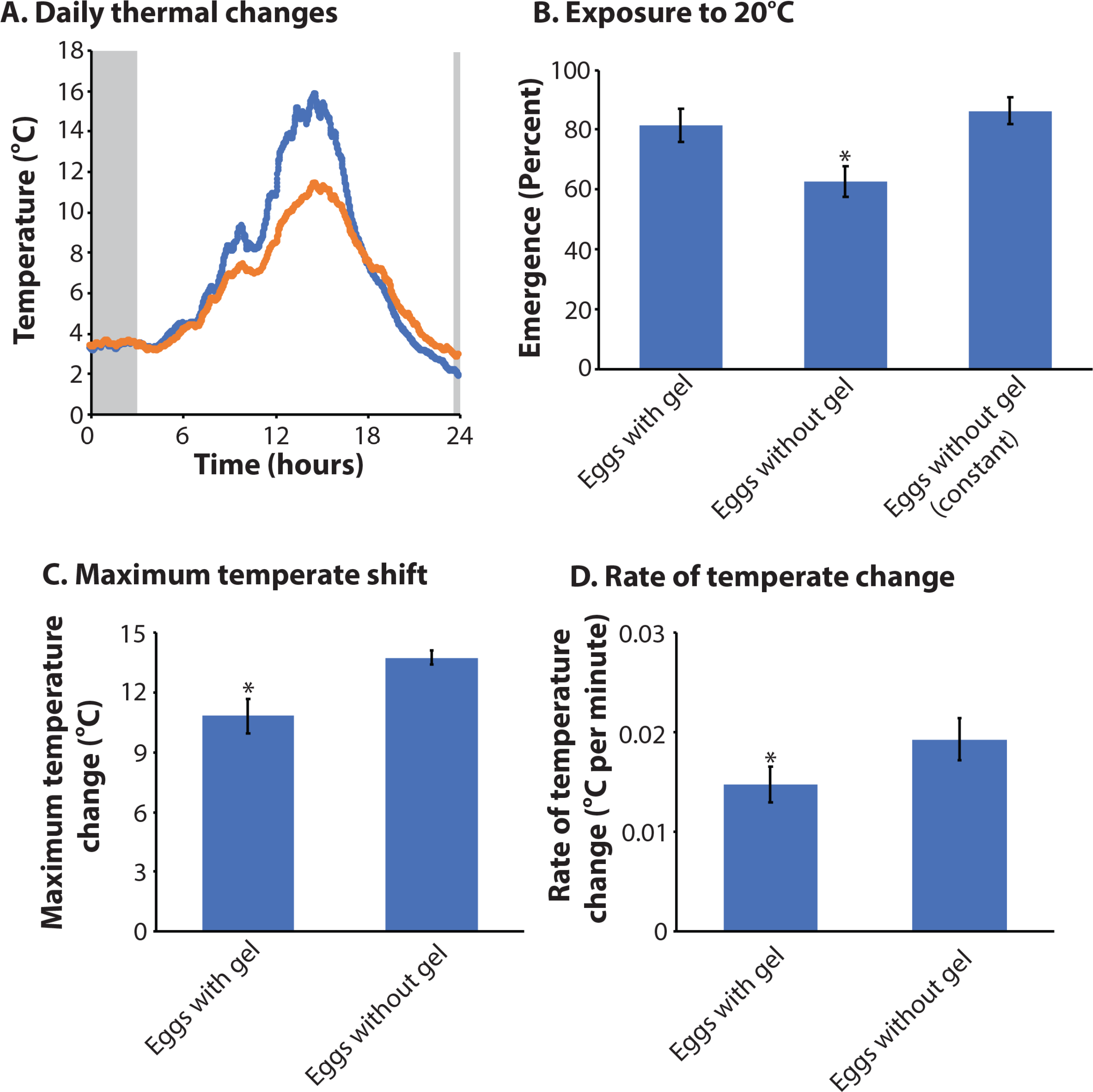
Role of accessory gland gel in relation to thermal buffering of eggs. A. Thermal profile within (orange) and outside (blue) gel-egg mixture under field conditions, B. Egg viability following exposure to 20℃ for three hours. Eggs without gel (constant) were held at 4°C for the duration of the trial. Maximum temperature change during 24 hour period. D. Rate of temperature change (minimum to maximum). Analysis of variance was utilized to examine statistical differences with the use of R statistic packages. * above indicates significance at P < 0.05 unless otherwise noted.

### 3.10 Population modeling

Dehydration of males, females, and both males and females together reduced population growth relative to control populations (Fig. 14). The combined effect of male and female dehydration had the largest effect, reducing population growth by nearly 21%. Lack of gel also negatively impact population growth, resulting in a 13% reduction relative to controls. The effect of thermal stress on egg viability would reduce population growth by 8%. In spite of these scenarios, populations would still be expected to increase in abundance under each specific stress. Under the “worst case scenario” where fecundity is reduced by male and female dehydration, larval survival is reduced by thermal stress, and there is a lack of gel reducing egg survival, population growth rate is negative (λ = -0.95), indicating declining population size. We note that all these growth rates are fairly liberal because of the relatively short time over which larval survival was observed (12 days) relative to the time spent as larvae in nature. In addition, it is likely that other factors, such as starvation, freezing, or pathogen attack, could occur, resulting in additive impacts on population growth.

**Figure 14:**
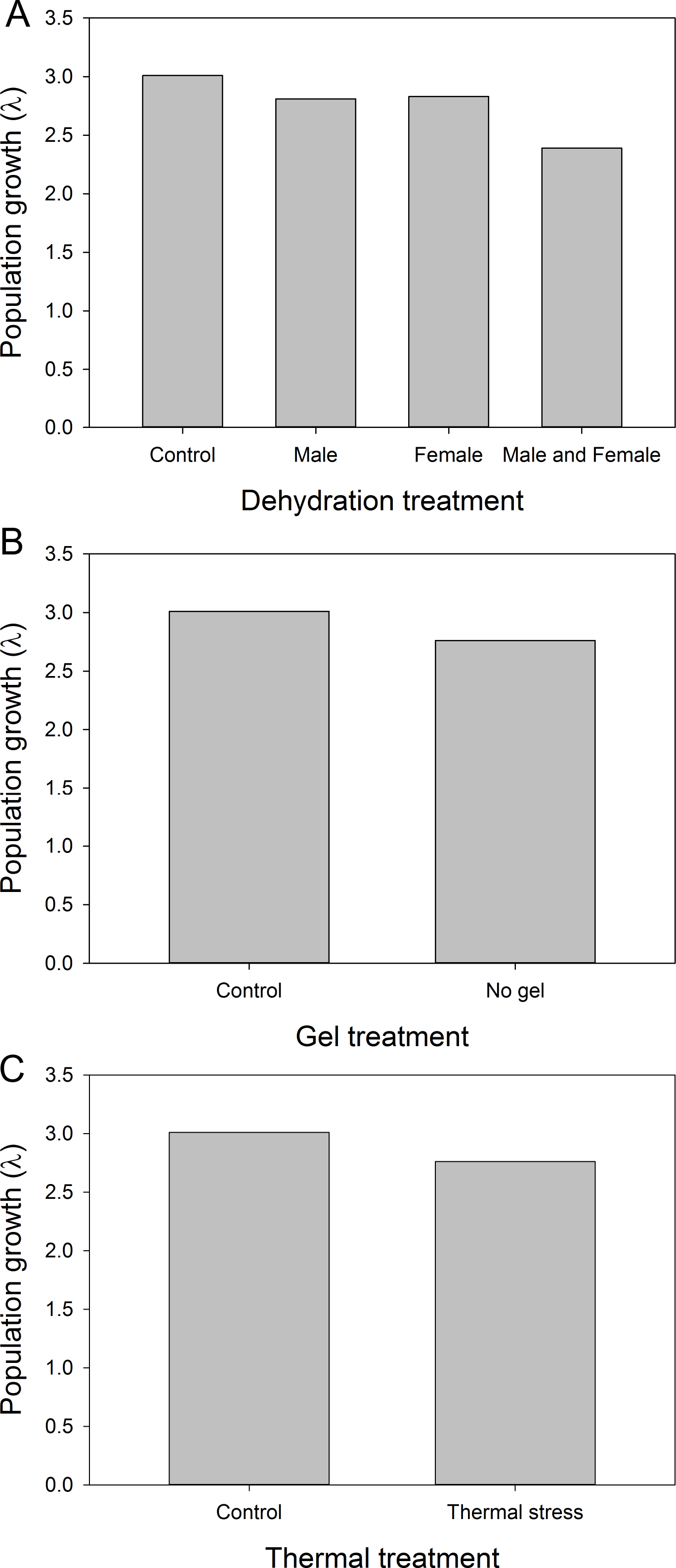
Population growth is impacted by dehydration and thermal stress in developing larvae. A. Population growth following altered egg production due to dehydration exposure as larvae in males, females, and both sexes combined compared to control (no dehydration of larvae). B. Growth based on the presence or absence of the gel under favorable conditions. C. Impact of thermal stress on egg viability with and without accessory gland gel.

## 4 DISCUSSION

This study examined reproductive biology of the Antarctic midge with the goal of establishing key molecular mechanisms associated with male and female biology. Combined RNA-seq and proteomics established the transcriptional components of reproduction and protein constituents deposited in the gel surrounding the egg. Specifically, we examined whether the gel that encases the eggs alters egg viability and larval survival and examined the impact of larval dehydration exposure on adult fertility. Little is known about midge reproductive biology and we have summarized the major findings of this study in Figure 15. Thus, this study will hopefully provide a foundation for the fields of Antarctic and midge reproductive biology. Lastly, we modeled the impact of multiple reproductive factors on population growth, each of which exert minor effects but in combination could yield a negative growth rate. Our results highlight the importance of understanding the reproductive biology of this Antarctic insect, a species restricted to a limited geographic region and a specific habitat.

**Figure 15:**
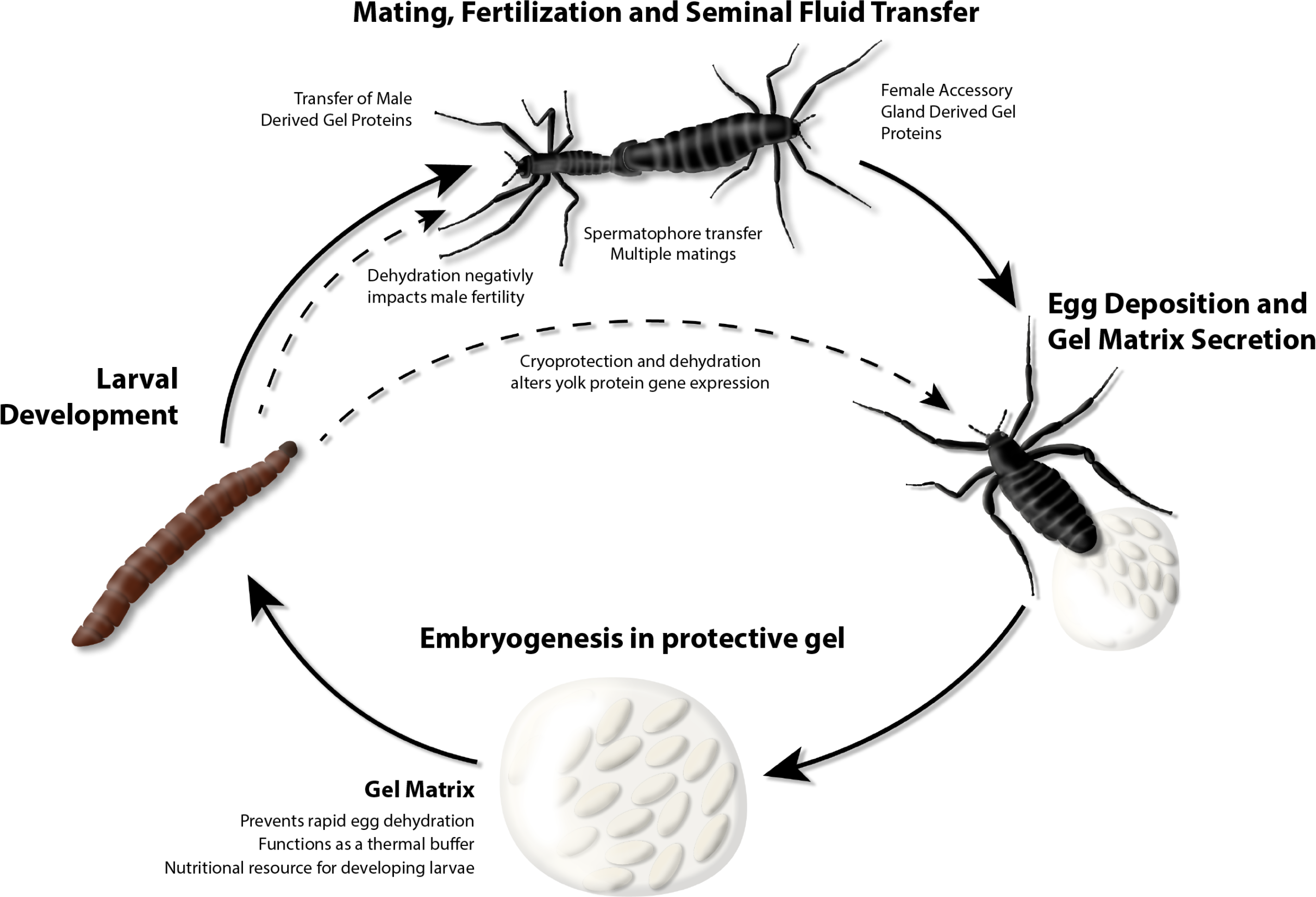
Summary of Antarctic midge reproduction. Larval development (four stages) is condensed into a single representation or all stages. Adults live approximately two-three weeks. Egg development occurs over thirty days. Impact of specific conditions are highlighted based on experimental evidence from this study.

The large proportion of apparently conserved, female-specific or female-enriched genes that are uncharacterized (≈44%) suggests there are many aspects of *Belgica* female physiology that remain poorly understood. However, the results of the GO analysis of the female-enriched gene set are consistent with results reported for *Drosophila* and other insect systems (Attardo et al., 2019; Olafson et al., 2019; Panfilio et al., 2019; Swanson, Wong, Wolfner, & Aquadro, 2004). In an analysis of genes differentially expressed between germline-naive and gonadectomized females in *Drosophila*, Parisi *et al*. found enrichment in terms associated with metabolism (Parisi et al., 2004). In fact, genes associated with energy storage and utilization, such as digestive proteases or lipid storage genes, showed increased expression in germline-naive males and females. This suggests that enrichment of such genes would be expected in the larval gene set, even if adult *Belgica* were active eaters which they are not. Indeed, the larva enriched gene set is replete with putative trypsins, chymotrypsin like proteins, and lipases (Cao & Jiang, 2017; Graveley et al., 2011; McKenna et al., 2016; Pauchet et al., 2010; Venancio, Cristofoletti, Ferreira, Verjovski-Almeida, & Terra, 2009). Oogenic gene expression has also been linked to nutrient sensing in *Drosophila* (Shim, Gururaja-Rao, & Banerjee, 2013; Terashima & Bownes, 2005). Enrichment of terms associated with metabolism, including components of ribosomes as protein-building machinery, identified in *B. antarctica* is most likely associated with oogenesis-related biosynthesis. This could include mobilizing lipids and proteins from the adult fat body during vitellogenesis, as well as synthesis of components of the egg gel excreted from the accessory gland. Transcription factor analyses revealed a subset of genes likely to have critical roles in female biology and thus warrant further more directed studies. One identified was mothers against decapentaplegic (Mad) homolog 1, which likely is likely critical for development (Newfeld, Chartoff, Graff, Melton, & Gelbart, 1996; Xie & Spradling, 1998).

Many enriched GO terms can also be tied to gametogenesis and embryogenesis or *Belgica* females. Terms associated with cell cycle control and DNA/RNA/chromatin handling are among the main examples. Beyond meiotic divisions that result in formation of gametes, which of course takes place in the male as well, female may have to exercise more precise control over the timing of oocyte maturation, deploying cell-cycle modulators, transcription regulation machinery, and/or RNA silencing factors (Lefebvre & Lécuyer, 2018; Qazi, Heifetz, & Wolfner, 2003; Soller, Bownes, & Kubli, 1999). TF analyses revealed a NF-kappa relish-like transcription factor as a potential regulator of transcript levels for the female-enriched gene set. Indeed, NF-kappaB transcription factors play well-established roles in transcriptional regulation of cytokines, molecules involved in cell cycle control, and development (Gilmore, 2006; Kim & Kim, 2005; Sosic & Olson, 2003). Curiously, this TF has its highest expression in males despite a paucity of binding sites in the regulatory regions of the male-enriched gene set. However, expression in the male accessory gland is limited and lower than that of the female or female accessory gland. The meaning of these findings remains unclear, but indicate a potential critical role related to accessory gland function in *Belgica*.

The female also produces developmental factors that play a role in successful embryogenesis, such as mRNAs, hormone biosynthesis precursors, and general cytoplasmic components (sperm are quite cytoplasm-poor relative to eggs) (Swevers, Raikhel, Sappington, Shirk, & Iatrou, 2005). Among the most highly enriched genes in females relative to males (>1000 fold higher) are a handful of genes known to be involved in embryonic patterning in insects, such as *nanos*, *oskar*, and multiple innexins, components of invertebrate gap junctions involved in intercellular communication during embryogenesis (Bauer et al., 2005; De Keuckelaere, Hulpiau, Saeys, Berx, & Van Roy, 2018; Güiza, Barria, Saez, & Vega, 2018; Quinlan, 2016; Richard & Hoch, 2015). This lends support to the notion that the female-enriched gene set is comprised largely of genes regulating and supporting oogenesis and embryogenesis. In addition, paternal DNA must be de-compacted after fertilization, meaning that sperm nucleotide binding proteins must be degraded (Doyen et al., 2015). The enzymes/proteins that participate in this process are likely already present in the egg, and therefore maternally contributed (Tirmarche, Kimura, Dubruille, Horard, & Loppin, 2016). This is followed by histone/chromatin assembly which is dependent on maternally provided histones and nucleosome assembly factors (Loppin, Dubruille, & Horard, 2015; Tirmarche et al., 2016). Male-contributed mitochondrial DNA must also be degraded (DeLuca & O’Farrell, 2012; Sato & Sato, 2011). This further accounts for the prevalence of terms associated with histones and chromosome organization. Finally, in *Drosophila,* nurse cells surrounding the oocyte are known to polyploidize regions of their nuclear DNA to enhance transcription and support provisioning of the oocyte (Bastock & St Johnston, 2008; Kaulenas, 1992; Orr-Weaver, 2015). This provides yet another physiological link to DNA replication and chromosome organization, but chironomids lack nurse cells. It is uncertain whether other cells of the ovary act in a similar manner to *Drosophila* nurse cells (Bastock & St Johnston, 2008).

The dominant GO terms associated with the male-enriched gene set are ‘anion transport’ and ‘carboxylic acid biosynthesis.’ These are both terms that can be linked to spermatogenesis and may also be linked to the differing metabolic needs of males, females, and larvae: larvae eat, digest, and grow; females conduct vitellogenesis, oogenesis, and embryogenesis; and males likely devote the bulk of their energy store to spermatogenesis and mating (Perry, Harrison, & Mank, 2014). The two terms may be interrelated. For example, genes involved in synthesis and transport of pyruvate, a carboxylate anion and the starting substrate of the tricarboxylic acid (TCA) cycle, could be associated with either GO term. The functional role of pyruvate metabolism in sperm viability is not known, but mitochondrial activity is demonstrated to be important for fertility, and a mitochondrial pyruvate transporter that is almost uniquely expressed in the male germline has been identified in placental mammals (Vanderperre et al., 2016). However, inhibition of pyruvate transporters in the mitochondrial membrane does not appear to affect sperm motility, lending support to the notion that sperm motility is fueled primarily by glycolysis (not import of pyruvate into mitochondria) (Vanderperre et al., 2016); alternatively, amino acid metabolism may play a role in motility, as it has been associated with sperm motility in *An. gambiae* (Izquierdo et al., 2019). TCA-based energy production must play a role in some other aspect of sperm viability (Vanderperre et al., 2016). The fact that a mitochondrial pyruvate transporter is important in sperm viability suggests that pyruvate transport and metabolism contribute to the significance of the GO terms ‘tricarboxylic acid cycle,’ ‘monocarboxylic acid metabolic process,’ and ‘anion transport.’ Additionally, sperm are thought to take up lactate and pyruvate through general monocarboxylic acid transporters (MCTs) in humans (Ramalho-Santos et al., 2009; Rato et al., 2012; Vanderperre et al., 2016). Perhaps there is an analogous uptake of energetically important organic acids in the seminiferous tubules of *Belgica* and other insects. Finally, pyruvate may be useful in supplying Acetyl-CoA for the histone acetylation that is essential for chromatin condensation during spermatogenesis (Ramalho-Santos et al., 2009; Rato et al., 2012; Vanderperre et al., 2016).

Overall, males showed far fewer genes with enriched expression relative to the female than vice versa. Of the 12 genes that are >1000 fold higher in the male compared to other stages, three are putative metalloendopeptidases. These may be involved in the proteolytic processing of amyloid precursor proteins (APPs), which are integral components of sperm membranes in humans, though their specific roles have not been determined (Silva et al., 2015). Also of interest in this small set of highly enriched genes are a few genes commonly, though not exclusively, associated with immunity: a leucine rich immune protein TM, a toll protein, and an apparent homolog of the transcription factor NF-X1 (Brucker, Funkhouser, Setia, Pauly, & Bordenstein, 2012; De Gregorio, Spellman, Rubin, & Lemaitre, 2001; Irving et al., 2001; Palmer & Jiggins, 2015). This transcription factor binds X-box motifs to regulate many eukaryotic genes and is generally thought to be a negative regulator of transcription (Stroumbakis, Li, & Tolias, 1996). The homolog of this transcription factor identified in *D. melanogaster*, named shuttlecraft (stc), is essential for normal embryonic development and is expressed most highly in the embryo CNS. Moreover, expression of this gene is highest in the ovary, and the promoters of many maternally contributed genes known to be crucial in embryonic development contain X-box motifs, such as *oskar, torpedo, pumilio,* and vitelline membrane proteins (Stroumbakis et al., 1996). Therefore it is tempting to suggest that, unlike in *Drosophila*, it is for some reason the male who produces this critical TF, transferring it to the female during copulation in order to trigger or facilitate proper development of their progeny. In support of the notion that the NF-X1 gene product is transferred to the female during copulation, expression of the gene is roughly 24-fold higher in the male accessory gland than in the whole carcass.

Gene ontology analysis of the larvae-enriched gene set revealed a preponderance of terms related to peptidase, hydrolase, and detoxification activity. These GO terms are related to digestion and detoxification of ingested materials by the growing larvae – neither of which would be relevant issues for the non-feeding adults. One cytochrome p450s (CYP6Z2), up-regulated in larvae, has been implicated in chemical resistance in *Anopheles* mosquitoes(Malik et al., 2016) and therefore may be critical for larval survival in their potentially toxic microhabitat, such as seabird guano (Rial et al., 2016).

Besides the ingestion of food, larvae of *B. antarctica* are also longer-lived and must face the seasonal challenges of permanent residence in Antarctica (Edwards, & Baust, 1981; Sugg et al., 1983). In particular, larvae must survive freezing and desiccation during the austral winter along with potential thermal stress during summer (Benoit et al., 2009; Benoit, Lopez-Martinez, Michaud, et al., 2007; Lopez-Martinez et al., 2008; Michaud et al., 2008; Rinehart et al., 2006; Teets, Kawarasaki, Lee, & Denlinger, 2013). Genes enhancing survival under these conditions, such as heat shock proteins (Benoit et al., 2009; Benoit, Lopez-Martinez, Michaud, et al., 2007; Lopez-Martinez et al., 2008; Michaud et al., 2008; Rinehart et al., 2006; Teets et al., 2013), are expected to be up-regulated by larvae. Based on previous studies in *Belgica* and other species that are desiccation and freeze tolerant, genes of importance could include oxido-reductase related enzymes (e.g. cytochrome p450s), Late Embryogenesis Abundant (LEA) proteins, protein repair methyltransferases, hemoglobins, aquaporins, and enzymes involved in trehalose metabolism (Benoit et al., 2009; Benoit, Lopez-Martinez, Michaud, et al., 2007; Dunning et al., 2013; Lopez-Martinez et al., 2008; Lv et al., 2010; Michaud et al., 2008; Rinehart et al., 2006; Ronges, Walsh, Sinclair, & Stillman, 2012; Teets et al., 2013; J. Zhang, Marshall, Westwood, Clark, & Sinclair, 2011). Many of these categories are indeed enriched in the larvae and are likely substantial factors in the stress resistance abilities of *B*. *antarctica*. Note that the presence of hemoglobin in insect hemolymph is unique to chironomids (Gusev et al., 2014; Kaiser et al., 2016; S.-M. Lee, Lee, Park, & Choi, 2006) and furthermore it is thought to be unique to the larval stage. This could partially explain the prevalence of iron binding as a GO term enriched in the larvae and conserved among the midge species analyzed (Gusev et al., 2014; Kaiser et al., 2016; Lee et al., 2006).

Most prevalent among GO terms enriched in the larval gene set are those associated with aminoglycan metabolism. This set includes, for example, multiple putative N-acetylgalactosaminyltransferases, which catalyze the initial glycosylation of serine and threonine residues (Tran & Ten Hagen, 2013). Mucin type O-glycosylation occurs commonly on proteins with extracellular domains, comprising a portion of the extracellular matrix, and it is thought to be important in intercellular communication and adhesion (Tran & Ten Hagen, 2013). In *Drosophila*, enzymes responsible for building glycosylated proteins, such as mucins, are critical for embryonic development, particularly for the CNS (Tian & Hagen, 2007; Tran & Ten Hagen, 2013; Zhang, Zhang, & Ten Hagen, 2008). Maternal and zygotic O-linked glycans have also been implicated in proper respiratory development. These O-glycans are termed *pgant* in *Drosophila* (Tian & Hagen, 2007; Tran & Ten Hagen, 2013; Zhang et al., 2008). One essential gene, *pgant4*, is involved in regulating gut acidification, but this is not the only such gene required in the digestive system (Tran & Ten Hagen, 2013; Tran et al., 2012). The larva enriched gene set in *Belgica* includes putative homologs of *pgant4* and *pgant6*, as well as the essential genes CG30463 and C1GalTA, both involved in mucin type O-linked glycosylation. However, larva-enriched GO terms, as mentioned above, are dominated by hydrolysis and catabolic processes, including glycosaminoglycan catabolism. This may be a sign of the breakdown of extracellular materials associated with growth and development or an indication of a diet rich in aminoglycans.

Composition of male accessory glands has been a major focus in numerous insects. These products influence fertilization rates, subsequent female receptivity to courtship, and are critical in the composition of the spermatophore (Alfonso-Parra et al., 2016; Avila et al., 2011; Dottorini et al., 2007; Gabrieli et al., 2014; Izquierdo et al., 2019; McGraw et al., 2008; Mitchell et al., 2015; Ravi Ram & Wolfner, 2007; Rogers et al., 2008; Sirot, Wong, Chapman, & Wolfner, 2015; Thailayil et al., 2011; Villarreal et al., 2018). Many of the male accessory gland genes from *Belgica* have orthologs based on predicted genes from midge and mosquito genomes, but few overlapping orthologs were identified that are expressed in the male accessory gland of *B. antarctica* compared to the male reproductive tissues of mosquitoes (Izquierdo et al., 2019; Papa et al., 2017). This is not surprising since a similar lack in orthology among male accessory gland products has been observed between *Drosophila* and *Glossina* and other higher flies (Attardo et al., 2019; Scolari et al., 2016). We did not conduct biological examination of specific roles for accessory gland proteins from *B. antarctica*, such as whether their transfer during mating impacts refractoriness of females, as noted in many species (Avila et al., 2011; Baldini, Gabrieli, Rogers, & Catteruccia, 2012; Ravi Ram & Wolfner, 2007; Sirot et al., 2015). There is an enrichment for genes associated with glycoprotein synthase, which is unsurprising as glycoproteins are common constituents of seminal fluid (Avila et al., 2011; Poiani, 2006). In addition, there were specific serine proteases, immune factors, and products involved in response to oxidative stress. These are similar to those observed in other fly species(Avila et al., 2011; Baldini et al., 2012; Ravi Ram & Wolfner, 2007; Sirot et al., 2015; Tian et al., 2017) and likely serve to preserve sperm viability but could impact other aspects such as female biology (Abraham et al., 2016; Denis et al., 2017). These factors are likely critical to male success, as this midge will mate on multiple occasions. It is important to note that females will deposit eggs with and without fertilization (Harada et al., 2014), thus suggesting that factors supplied in the spermatophore are not essential for ovulation and oviposition as in other fly species (Avila et al., 2011; Baldini et al., 2012; Ravi Ram & Wolfner, 2007; Sirot et al., 2015).

Our gene expression analysis is one of the few that has explicitly examined gene expression within the female accessory gland. Many, but not all, products present within the gel were expressed in the female’s accessory gland. A comparison of our results to expression pattern in mosquito female tracts (Papa et al., 2017) revealed few genes with overlapping expression(Papa et al., 2017). Few studies in other flies explicitly examined products generated by the female’s accessory gland. Even in *Drosophila*, the female accessory gland (paraovaria) remains one of the most understudied organs (Attardo et al., 2014). In tsetse flies, this organ provides nourishment to developing intrauterine larvae (Attardo et al., 2019; Benoit et al., 2015; Attardo et al., 2014), but there are no similarities, beyond standard housekeeping genes, between this organ in tsetse flies and *B. antarctica*. Of interest, regulation of transcript expression within the female accessory gland seems to be conserved (Attardo et al., 2014), suggesting that regulatory aspects may be similar but drive expression of specific genes related to unique functions of this organ in different flies. Transcription factor analyses identified a single gene (IU25_08656) that has enriched levels of both binding sites upstream of female accessory gland enriched genes and is itself expressed in the female accessory gland. This is an uncharacterized zinc finger proteins, but is likely to have a critical role in female reproductive function for *B*. *antarctica*.

The three major protein components of the gel are vitellogenin, larval serum protein, and apolipophorin, all which are reasonable components of a gel whose primary function includes fueling the development of larvae upon hatching. *Belgica* larvae exhibit “drinking behavior” shortly before hatching, suggesting that up to this point embryonic development is fueled by nutrient reserves present in the egg at the time of oviposition (Harada et al., 2014). Upon hatching, the gel is ingested by the larva, making it the first meal fueling their development. Larval serum protein likely acts as an amino acid reservoir, while vitellogenin and apolipoproteins could provide fatty acid, sugar, and protein reserves. The gel may also contribute other elements in smaller amounts, such as pre-synthesized developmental hormones in an inactive form, hormone precursors, proenzymes, and enzyme cofactors that are important for continued development (Harada et al., 2014). Larval serum protein is a storage protein belonging to the family of hexamerins (Burmester, 1999; Burmester et al., 1998; Telfer & Kunkel, 1991). Genes for such proteins are often highly expressed in the final larval instar preceding pupation (Burmester, 1999; Burmester et al., 1998; Telfer & Kunkel, 1991) and serve as a nutrient reserve for developing pupae and newly emerged adult. During the sweeping morphological changes that accompany metamorphosis in holometabolous insects like *Belgica*, such storage proteins are extracted from the larval fat body, transferred into the hemolymph, and subsequently re-sequestered in the newly formed adult fat body(Burmester, 1999; Burmester et al., 1998; Keeley, 1978; Larsen, 1976; Telfer & Kunkel, 1991). The following scenario is the likely fate of the larval serum protein found in the egg gel: Initially re-sequestered by the adult fat body and shuttled to the female accessory gland with vitellogenin, or possibly sequestered in the female accessory gland during adult development after pupation without a layover in the fat body. Importantly, lack of expression in the female adult indicates these proteins must be synthesized in the larva for subsequent use by the adult.

Along with establishing components of the female accessory gland derived gel, a major goal of this study was to identify functional roles of this gel and to determine if the gel components are impacted by stress in the developing larvae. Based on our results, one of the main functions of the gel is to provide a nutritional resource, which leads to a higher larval survival. Products of the accessory glands of females have been documented as food sources for insect species among many orders(Attardo et al., 2019; Benoit et al., 2015; Denlinger & Ma, 1974; Kaiwa et al., 2014; Ma, Denlinger, Jarlfors, & Smith, 1975). Production of gel substances surrounding eggs have been noted in other species, such as in stink bugs, where the gel serves as a source of nutrition, protects the eggs, and provides a vehicle to transfer microbial symbionts (Kaiwa et al., 2014). Similar to our study, removal of the gel inhibits juvenile development in stink bugs, but in contrast, removal of the gel around the stink bug eggs has little impact on hatching success. In addition to providing a critical nutritional source and preventing dehydration, we show that the egg gel acts as a thermal buffer that limits temperature extremes. Our *de novo* assembly of the female accessory gland failed to detect signatures of microbial symbionts within the gel, indicating that the gel does not serve as a mechanism of bacterial transfer to the developing larvae. This lack of microbial presence could very well limit microbial exposure of the midge until after larvae emerge from the gel. Accessory gland products also have been demonstrated to possess antimicrobial properties(Avila et al., 2011; Otti, McTighe, & Reinhardt, 2013; Otti, Naylor, Siva-Jothy, & Reinhardt, 2009); this is a possibility for *B. antarctica* because immune peptides are present in the gel.

Larval dehydration stress had a major impact on fertility of both adult males and females. The most likely cause of reduced fecundity in males and females is a direct reduction in larval serum protein, a hexamerin that acts as a storage protein in larvae (Burmester, 1999; Burmester et al., 1998; Telfer & Kunkel, 1991). This hexamerin represents one of the highest expressed transcripts in developing larvae, so its accumulation during the juvenile stages is likely a critical resource for production of eggs and accessory gland components. This effect is likely even more pronounced in *B. antarctica* because adults do not feed or even readily drink water (Benoit, Lopez-Martinez, Elnitsky, Lee, & Denlinger, 2007). Along with acting as a nutritional source, the larval serum protein generated in juvenile stage of females is likely incorporated into the accessory gland gel, suggesting that a reduction in this product may directly impact gel composition. Larval nutritional status has a direct impact on fecundity in numerous insect systems (Aguila, Hoshizaki, & Gibbs, 2013; Aguila, Suszko, Gibbs, & Hoshizaki, 2007; Rosa & Saastamoinen, 2017), including other midges (Sibley et al., 2001). To the best of our knowledge, larval dehydration has not previously been examined in relation to subsequent adult fecundity. Other stressful conditions, such as chemical exposure, have impaired both larval development and subsequent adult reproduction in a non-biting midge, *Chironomus riparius*(Vogt et al., 2007).

This study provides an encompassing view of reproductive biology of the Antarctic midge, from molecular mechanisms to the impact of larval stress exposure on adult fecundity. Key findings are summarized in Figure 15. This is followed by population growth modeling to establish how these factors directly impact persistence of this insect in its limited Antarctic range. Population modeling revealed that each factor (dehydration stress, lack of gel, thermal buffering), by itself, has a small impact on population growth but combined factors likely result in negative population growth. The limited reproductive window of 2-3 weeks makes understanding both male and female reproduction critical for understanding how this midge survives in Antarctica. Studies on the reproductive biology of flies have been limited largely to *Drosophila* and disease vectors (sand flies, mosquitoes, etc.), and our results expand into Chironomidae to provide the groundwork for future studies with this dipteran system.

## ACKNOWLEDGEMENTS

This work was supported by the National Science Foundation grant DEB-1654417 (partially) and United States Department of Agriculture 2018-67013 (partially) to J.B.B., National Science Foundation grant OPP-1341393 to D.L.D., and National Science Foundation grant OPP-1341385 to R.E.L. We thank the staff at Palmer Station, Antarctica for assistance in logistics, experiments, and helping make the field season a pleasant experience.

## AUTHOR CONTRIBUTION

This study was designed and conceived by J.B.B. The manuscript was written by G.F. and J.B.B. RNA-seq studies were conducted by G.F., A.J.R., B. D., and S.T.B. Molecular and physiological studies were conducted and interpreted by S. N., C. P., G. F., C.J.H, E.M.D., and J.B.B. Transcription factor analyses were contributed by X.C. and M.T.W. and interpreted by J.B.B. and G.F. Additional data and statistical analyses were conducted by K.J.O., G.M.A., and J.B.B. Samples were collected by J.D.G., D.S., R.E.L., and D.L.D. Population modeling was conducted by S.F.M. Figures were prepared by G.F. and J.B.B. Illustrations were by G.M.A. All authors were responsible for editing the manuscript and have approved publication.

## DATA AVAILABILITY STATEMENT

All data generated for this project have been submitted to NCBI or made available in the supplemental files.

**Supplemental table 1.** RNA-seq results for complete *Belgica antarctica* gene set. Expression values are in transcripts per million. RNA-seq datasets are available under the following NCBI Bioprojects PRJNA174315 and PRJNA576639.

**Supplemental table 2**-BLAST results for female accessory gland specific *de novo* transcriptome against bacterial sequences from the NCBI nr database.

**Supplemental table 3.** RNA-seq results for female-enriched *Belgica antarctica* gene set. Expression values are in transcripts per million. RNA-seq datasets are available under the following NCBI Bioprojects PRJNA174315 and PRJNA576639.

**Supplemental table 4.** RNA-seq results for male-enriched *Belgica antarctica* gene set. Expression values are in transcripts per million. RNA-seq datasets are available under the following NCBI Bioprojects PRJNA174315 and PRJNA576639.

**Supplemental table 5.** RNA-seq results for larvae-enriched *Belgica antarctica* gene set. Expression values are in transcripts per million. RNA-seq datasets are available under the following NCBI Bioprojects PRJNA174315 and PRJNA576639.

**Supplemental table 6.** RNA-seq results for female accessory gland-enriched *Belgica antarctica* gene set. Expression values are in transcripts per million. RNA-seq datasets are available under the following NCBI Bioprojects PRJNA174315 and PRJNA576639.

**Supplemental table 7.** RNA-seq results for male accessory gland-enriched *Belgica antarctica* gene set. Expression values are in transcripts per million. RNA-seq datasets are available under the following NCBI Bioprojects PRJNA174315 and PRJNA576639.

**Supplemental table 8.** WGCNA module results for female-enriched *Belgica antarctica* gene set. Expression values are in transcripts per million. RNA-seq datasets are available under the following NCBI Bioproject PRJNA174315 and PRJNA576639.

**Supplemental table 9.** WGCNA module results for male-enriched *Belgica antarctica* gene set. Expression values are in transcripts per million. RNA-seq datasets are available under the following NCBI Bioprojects PRJNA174315 and PRJNA576639.

**Supplemental table 10.** WGCNA module results for larvae-enriched *Belgica antarctica* gene set. Expression values are in transcripts per million. RNA-seq datasets are available under the following NCBI Bioproject PRJNA174315 and PRJNA576639.

**Supplemental table 11.** WGCNA module results for female accessory gland-enriched *Belgica antarctica* gene set. Expression values are in transcripts per million. RNA-seq datasets are available under the following NCBI Bioproject PRJNA174315 and PRJNA576639.

**Supplemental table 12.** WGCNA module results for male accessory gland-enriched *Belgica antarctica* gene set. Expression values are in transcripts per million. RNA-seq datasets are available under the following NCBI Bioproject PRJNA174315 and PRJNA576639.

**Supplemental table 13.** Transcript levels for gel-derived proteins from the *Belgica antarctica* gene set. Expression values are in transcripts per million. RNA-seq datasets are available under the following NCBI Bioproject PRJNA174315 and PRJNA576639. Relative abundance of protein amounts were based on the number of protein fragments that match a specific genes.

**Supplemental table 14.** Transcript levels for transcription factors from the *Belgica antarctica* gene set. Expression values are in transcript per million. RNA-seq datasets are available under the following NCBI Bioproject PRJNA174315 and PRJNA576639.

**Supplemental table 15-** Quantitative PCR primers used for the validation of RNA-seq result.

## REFERENCES

1. Abbott, K. C., & Dwyer, G. (2007). Food limitation and insect outbreaks: complex dynamics in plant–herbivore models. Journal of Animal Ecology, 76(5), 1004–1014.

2. Abraham, S., Lara-Pérez, L. A., Rodríguez, C., Contreras-Navarro, Y., Nuñez-Beverido, N., Ovruski, S., & Pérez-Staples, D. (2016). The male ejaculate as inhibitor of female remating in two tephritid flies. Journal of Insect Physiology, 88, 40–47.

3. Afgan, E., Baker, D., Batut, B., Van Den Beek, M., Bouvier, D., Čech, M., … Grüning, B. A. (2018). The Galaxy platform for accessible, reproducible and collaborative biomedical analyses: 2018 update. Nucleic acids research, 46(W1), W537–W544.

4. Aguila, J. R., Hoshizaki, D. K., & Gibbs, A. G. (2013). Contribution of larval nutrition to adult reproduction in *Drosophila melanogaster*. Journal of Experimental Biology, 216(3), 399–406.

5. Aguila, J. R., Suszko, J., Gibbs, A. G., & Hoshizaki, D. K. (2007). The role of larval fat cells in adult *Drosophila melanogaster*. Journal of Experimental Biology, 210(6), 956–963.

6. Alfonso-Parra, C., Ahmed-Braimah, Y. H., Degner, E. C., Avila, F. W., Villarreal, S. M., Pleiss, J. A., … Harrington, L. C. (2016). Mating-induced transcriptome changes in the reproductive tract of female *Aedes aegypti*. PLoS Neglected Tropical Diseases, 10(2), e0004451.

7. Attardo, G. M., Abd-Alla, A. M., Acosta-Serrano, A., Allen, J. E., Bateta, R., Benoit, J., … Christensen, M. B. (2019). The Glossina genome cluster: comparative genomic analysis of the vectors of African trypanosomes. bioRxiv, 531749.

8. Attardo, G. M., Benoit, J. B., Michalkova, V., Patrick, K. R., Krause, T. B., & Aksoy, S. (2014). Ladybird late homeodomain factor regulates lactation specific expression of milk proteins during pregnancy in the tsetse fly (*Glossina morsitans*). PLoS neglected tropical diseases, 8, e2645.

9. Avila, F. W., Sirot, L. K., LaFlamme, B. A., Rubinstein, C. D., & Wolfner, M. F. (2011). Insect seminal fluid proteins: identification and function. Annual review of entomology, 56, 21–40.

10. Baldini, F., Gabrieli, P., Rogers, D. W., & Catteruccia, F. (2012). Function and composition of male accessory gland secretions in *Anopheles gambiae*: a comparison with other insect vectors of infectious diseases. Pathogens and global health, 106(2), 82–93.

11. Bastock, R., & St Johnston, D. (2008). *Drosophila* oogenesis. Current Biology, 18(23), R1082–R1087.

12. Bauer, R., Löer, B., Ostrowski, K., Martini, J., Weimbs, A., Lechner, H., & Hoch, M. (2005). Intercellular communication: the *Drosophila* innexin multiprotein family of gap junction proteins. Chemistry & biology, 12(5), 515–526.

13. Benoit, J., Kölliker, M., & Attardo, G. (2019). Putting invertebrate lactation in context. *Science (New York*, NY), 363(6427), 593–593.

14. Benoit, J. B., Adelman, Z. N., Reinhardt, K., Dolan, A., Poelchau, M., Jennings, E. C., … Richards, S. (2016). Unique features of a global human ectoparasite identified through sequencing of the bed bug genome. Nature Communications, 7. doi:10.1038/ncomms10165

15. Benoit, J. B., Attardo, G. M., Baumann., A. A., Michalkova, V., & Aksoy, S. (2015). Adenotrophic viviparity in tsetse flies: potential for population control and as an insect model for lactation. Annual review of entomology, In press

16. Benoit, J. B., Lopez-Martinez, G., Elnitsky, M. A., Lee, R. E., & Denlinger, D. L. (2007). Moist habitats are essential for adults of the Antarctic midge, *Belgica antarctica* (Diptera : Chironomidae), to avoid dehydration. European Journal of Entomology, 104(1), 9–14.

17. Benoit, J. B., Lopez-Martinez, G., Elnitsky, M. A., Lee, R. E., & Denlinger, D. L. (2009). Dehydration-induced cross tolerance of *Belgica antarctica* larvae to cold and heat is facilitated by trehalose accumulation. Comparative Biochemistry and Physiology a-Molecular & Integrative Physiology, 152(4), 518–523. doi:DOI 10.1016/j.cbpa.2008.12.009

18. Benoit, J. B., Lopez-Martinez, G., Michaud, M. R., Elnitsky, M. A., Lee, R. E., Jr., & Denlinger, D. L. (2007). Mechanisms to reduce dehydration stress in larvae of the Antarctic midge, Belgica antarctica. Journal Insect Physiol, 53(7), 656–667. doi:10.1016/j.jinsphys.2007.04.006

19. Benoit, J. B., Michalkova, V., Didion, E. M., Xiao, Y., Baumann, A. A., Attardo, G. M., & Aksoy, S. (2018). Rapid autophagic regression of the milk gland during involution is critical for maximizing tsetse viviparous reproductive output. PLoS neglected tropical diseases, 12(1), e0006204.

20. Brucker, R. M., Funkhouser, L. J., Setia, S., Pauly, R., & Bordenstein, S. R. (2012). Insect Innate Immunity Database (IIID): an annotation tool for identifying immune genes in insect genomes. PLoS One, 7(9), e45125.

21. Burmester, T. (1999). Evolution and function of the insect hexamerins. European Journal of Entomology, 96, 213–226.

22. Burmester, T., Massey Jr, H. C., Zakharkin, S. O., & Benes, H. (1998). The evolution of hexamerins and the phylogeny of insects. Journal of Molecular Evolution, 47(1), 93–108.

23. Cao, X., & Jiang, H. (2017). An analysis of 67 RNA-seq datasets from various tissues at different stages of a model insect, *Manduca sexta*. BMC genomics, 18(1), 796.

24. Clark, A. G., Eisen, M. B., Smith, D. R., Bergman, C. M., Oliver, B., Markow, T. A., … MacCallum, I. (2007). Evolution of genes and genomes on the *Drosophila* phylogeny. Nature, 450(7167), 203–218. doi:nature06341 [pii]10.1038/nature06341

25. Clark, N., & Ma’ayan, A. (2011). Introduction to statistical methods for analyzing large data sets: Gene Set Enrichment Analysis (GSEA). Science Signaling, 4(190), tr4.

26. Conesa, A., Gotz, S., Garcia-Gomez, J. M., Terol, J., Talon, M., & Robles, M. (2005). Blast2GO: a universal tool for annotation, visualization and analysis in functional genomics research. Bioinformatics, 21(18), 3674–3676. doi:10.1093/bioinformatics/bti610

27. Convey, P. (1992). Aspects of the biology of the midge, *Eretmoptera murphyi* Schaeffer (Diptera: Chironomidae), introduced to Signy Island, maritime Antarctic. Polar Biology, 12(6-7), 653–657.

28. Convey, P. (1997). How are the life history strategies of Antarctic terrestrial invertebrates influenced by extreme environmental conditions? Journal of Thermal Biology, 22(6), 429–440.

29. Convey, P., & Block, W. (1996). Antarctic Diptera: ecology, physiology and distribution. European Journal of Entomology, 93, 1–14.

30. De Gregorio, E., Spellman, P. T., Rubin, G. M., & Lemaitre, B. (2001). Genome-wide analysis of the *Drosophila* immune response by using oligonucleotide microarrays. Proceedings of the National Academy of Sciences, 98(22), 12590–12595.

31. De Keuckelaere, E., Hulpiau, P., Saeys, Y., Berx, G., & Van Roy, F. (2018). Nanos genes and their role in development and beyond. Cellular and Molecular Life Sciences, 75(11), 1929–1946.

32. Degrugillier, M. E. (1985). In vitro release of house fly, *Musca domestica* L.(Diptera: Muscidae), acrosomal material after treatments with secretion of female accessory gland and micropyle cap substance. International Journal of Insect Morphology and Embryology, 14(6), 381–391.

33. DeLuca, S. Z., & O’Farrell, P. H. (2012). Barriers to male transmission of mitochondrial DNA in sperm development. Developmental cell, 22(3), 660–668.

34. Denis, B., Claisse, G., Le Rouzic, A., Wicker-Thomas, C., Lepennetier, G., & Joly, D. (2017). Male accessory gland proteins affect differentially female sexual receptivity and remating in closely related *Drosophila* species. Journal of Insect Physiology, 99, 67–77.

35. Denlinger, D. L., & Ma, W.-C. (1974). Dynamics of the pregnancy cycle in the tsetse Glossina morsitans Journal of insect physiology, 20, 1015–1026.

36. Dixon, S. M., Coyne, J. A., & Noor, M. A. (2003). The evolution of conspecific sperm precedence in *Drosophila*. Molecular Ecology, 12(5), 1179–1184.

37. Dottorini, T., Nicolaides, L., Ranson, H., Rogers, D. W., Crisanti, A., & Catteruccia, F. (2007). A genome-wide analysis in *Anopheles gambiae* mosquitoes reveals 46 male accessory gland genes, possible modulators of female behavior. Proceedings of the National Academy of Sciences, 104(41), 16215–16220.

38. Doyen, C. M., Chalkley, G. E., Voets, O., Bezstarosti, K., Demmers, J. A., Moshkin, Y. M., & Verrijzer, C. P. (2015). A testis-specific chaperone and the chromatin remodeler ISWI mediate repackaging of the paternal genome. Cell reports, 13(7), 1310–1318.

39. Dunning, L. T., Dennis, A. B., Park, D., Sinclair, B. J., Newcomb, R. D., & Buckley, T. R. (2013). Identification of cold-responsive genes in a New Zealand alpine stick insect using RNA-Seq. Comparative Biochemistry and Physiology Part D: Genomics and Proteomics, 8(1), 24–31. doi:10.1016/j.cbd.2012.10.005

40. Eddy, S. R. (2009). A new generation of homology search tools based on probabilistic inference. Genome Informatics 2009: Genome Informatics Series, 23, 205–211.

41. J.Edwards, J.S., & Baust, J. (1981). Sex ratio and adult behaviour of the Antarctic midge *Belgica antarctica* (Diptera, Chironomdae). Ecological Entomology, 6(3), 239–243.

42. Gabrieli, P., Kakani, E. G., Mitchell, S. N., Mameli, E., Want, E. J., Anton, A. M., … Catteruccia, F. (2014). Sexual transfer of the steroid hormone 20E induces the postmating switch in *Anopheles gambiae*. Proceedings of the National Academy of Sciences, 111(46), 16353–16358.

43. Giglioli, M., & Mason, G. (1966). The mating plug in anopheline mosquitoes. Paper presented at the Proceedings of the Royal Entomological Society of London. Series A, General Entomology.

44. Gilmore, T. D. (2006). Introduction to NF-κB: players, pathways, perspectives. Oncogene, 25(51), 6680.

45. Goecks, J., Nekrutenko, A., & Taylor, J. (2010). Galaxy: a comprehensive approach for supporting accessible, reproducible, and transparent computational research in the life sciences. Genome Biology, 11(8), R86.

46. Grabherr, M. G., Haas, B. J., Yassour, M., Levin, J. Z., Thompson, D. A., Amit, I., … Regev, A. (2011). Full - length transcriptome assembly from RNA-Seq data without a reference genome. Nature biotechnology, 29(7), 644–652. doi:nbt.1883 [pii]10.1038/nbt.1883

47. Graveley, B. R., Brooks, A. N., Carlson, J. W., Duff, M. O., Landolin, J. M., Yang, L., … Booth, B. W. (2011). The developmental transcriptome of *Drosophila melanogaster*. Nature, 471(7339), 473–479.

48. Güiza, J., Barria, I., Saez, J. C., & Vega, J. L. (2018). Innexins: Expression, Regulation and Functions. Frontiers in physiology, 9, 1414.

49. Gusev, O., Suetsugu, Y., Cornette, R., Kawashima, T., Logacheva, M. D., Kondrashov, A. S., … Shimura, S. (2014). Comparative genome sequencing reveals genomic signature of extreme desiccation tolerance in the anhydrobiotic midge. Nature Communication, 5, 4784.

50. Gwynne, D.T. (2012). Male mating effort, confidence of paternity, and insect sperm competition. Sperm competition and the evolution of animal mating systems, 117.

51. Hagan, R. W., Didion, E. M., Rosselot, A. E., Holmes, C. J., Siler, S. C., Rosendale, A. J., … Nine, G. A. (2018). Dehydration prompts increased activity and blood feeding by mosquitoes. Scientific reports, 8(1), 6804.

52. Hahn, S., & Reinhardt, K. (2006). Habitat preference and reproductive traits in the Antarctic midge *Parochlus steinenii* (Diptera: Chironomidae). Antarctic Science, 18(2), 175–181.

53. Harada, E., Lee, R. E., Denlinger, D. L., & Goto, S. G. (2014). Life history traits of adults and embryos of the Antarctic midge *Belgica antarctica*. Polar Biology, 37(8), 1213–1217.

54. Heaven, M. R., Funk, A. J., Cobbs, A. L., Haffey, W. D., Norris, J. L., McCullumsmith, R. E., & Greis, K. D. (2016). Systematic evaluation of data-independent acquisition for sensitive and reproducible proteomics—a prototype design for a single injection assay. Journal of Mass Spectrometry, 51(1), 1–11.

55. Hopkins, B. R., Sepil, I., Thézénas, M.-L., Craig, J. F., Miller, T., Charles, P. D., … Pizzari, T. (2019). Divergent allocation of sperm and the seminal proteome along a competition gradient in *Drosophila melanogaster*. Proceedings of the National Academy of Sciences, 116(36), 17925–17933.

56. Hopkins, B. R., Sepil, I., & Wigby, S. (2017). Seminal fluid. Current Biology, 27(11), R404–R405.

57. Huang da, W., Sherman, B. T., Zheng, X., Yang, J., Imamichi, T., Stephens, R., & Lempicki, R. A. (2009). Extracting biological meaning from large gene lists with DAVID. *Current Protocols in Bioinformatics*, Chapter 13. doi:10.1002/0471250953.bi1311s27

58. Initiative, I. G. G. (2014). Genome sequence of the tsetse fly (*Glossina morsitans*): vector of African trypanosomiasis. Science, 25(6182), 380–386.

59. Irving, P., Troxler, L., Heuer, T. S., Belvin, M., Kopczynski, C., Reichhart, J. M., … Hetru, C. (2001). A genome-wide analysis of immune responses in *Drosophila*. Proceedings of the National Academy of Sciences of the United States of America, 98(26), 15119–15124. doi:10.1073/pnas.261573998

60. Izquierdo, A., Fahrenberger, M., Persampieri, T., Benedict, M. Q., Giles, T., Catteruccia, F., … Dottorini, T. (2019). Evolution of gene expression levels in the male reproductive organs of *Anopheles* mosquitoes. Life science alliance, 2(1), e201800191.

61. Kaiser, T. S., Poehn, B., Szkiba, D., Preussner, M., Sedlazeck, F. J., Zrim, A., … Hummel, T. (2016). The genomic basis of circadian and circalunar timing adaptations in a midge. Nature, 540(7631), 69.

62. Kaiwa, N., Hosokawa, T., Nikoh, N., Tanahashi, M., Moriyama, M., Meng, X.-Y., … Ito, M. (2014). Symbiont-supplemented maternal investment underpinning host’s ecological adaptation. Current Biology, 24(20), 2465–2470.

63. Kaulenas, M. (1992). Structure and function of the female accessory reproductive systems. In Insect Accessory Reproductive Structures (pp. 33–121): Springer.

64. Kaulenas, M. (2012). Insect accessory reproductive structures: function, structure, and development (Vol. 31): Springer Science & Business Media.

65. Keeley, L. (1978). Endocrine regulation of fat body development and function. Annual Review of Entomology, 23(1), 329–352.

66. Kelley, J. L., Peyton, J. T., Fiston-Lavier, A.-S., Teets, N. M., Yee, M.-C., Johnston, J. S., … Denlinger, D. L. (2014). Compact genome of the Antarctic midge is likely an adaptation to an extreme environment. Nature Communications, 5, 4611.

67. Kennedy, A. D. (1993). Water as a limiting factor in the Antarctic terrestrial environment: a biogeographical synthesis. Arctic and Alpine Research, 25(4), 308–315.

68. Kim, S., Oh, M., Jung, W., Park, J., Choi, H.-G., & Shin, S. C. (2017). Genome sequencing of the winged midge, *Parochlus steinenii*, from the Antarctic Peninsula. GigaScience, 6(3), giw009.

69. Kim, T., & Kim, Y. (2005). Overview of innate immunity in *Drosophila*. Journal of biochemistry and molecular biology, 38(2), 121.

70. Kotrba, M. (1996). Sperm transfer by spermatophore in Diptera: new results from the Diopsidae. Zoological Journal of the Linnean Society, 117(3), 305–323.

71. Langfelder, P., & Horvath, S. (2008). WGCNA: an R package for weighted correlation network analysis. BMC bioinformatics, 9(1), 559.

72. Larsen, W. J. (1976). Cell remodeling in the fat body of an insect. Tissue and Cell, 8(1), 73–92.

73. Lee, K. P., Simpson, S. J., Clissold, F. J., Brooks, R., Ballard, J. W. O., Taylor, P. W., … Raubenheimer, D. (2008). Lifespan and reproduction in *Drosophila*: new insights from nutritional geometry. Proceedings of the National Academy of Sciences, 105(7), 2498–2503.

74. Lee, S.-M., Lee, S.-B., Park, C.-H., & Choi, J. (2006). Expression of heat shock protein and hemoglobin genes in *Chironomus tentans* (Diptera, chironomidae) larvae exposed to various environmental pollutants: a potential biomarker of freshwater monitoring. Chemosphere, 65(6), 1074–1081.

75. Lefebvre, F. A., & Lécuyer, É. (2018). Flying the RNA nest: *Drosophila* reveals novel insights into the transcriptome dynamics of early development. Journal of developmental biology, 6(1), 5.

76. Lefkovitch, L. (1965). The study of population growth in organisms grouped by stages. Biometrics, 1–18.

77. Leopold, R. A., & Degrugillier, M. E. (1973). Sperm penetration of housefly eggs: evidence for involvement of a female accessory secretion. Science, 181(4099), 555–557.

78. Lococo, D., & Huebner, E. (1980). The ultrastructure of the female accessory gland, the cement gland, in the insect *Rhodnius prolixus*. Tissue and Cell, 12(3), 557–580.

79. Lopez-Martinez, G., Benoit, J. B., Rinehart, J. P., Elnitsky, M. A., Lee, R. E., & Denlinger, D. L. (2009). Dehydration, rehydration, and overhydration alter patterns of gene expression in the Antarctic midge, *Belgica antarctica*. Journal of Comparative Physiology B-Biochemical Systemic and Environmental Physiology, 179(4), 481–491. doi:DOI 10.1007/s00360-008-0334-0

80. Lopez-Martinez, G., Elnitsky, M. A., Benoit, J. B., Lee, R. E., Jr., & Denlinger, D. L. (2008). High resistance to oxidative damage in the Antarctic midge *Belgica antarctica*, and developmentally linked expression of genes encoding superoxide dismutase, catalase and heat shock proteins. Insect Biochem Mol Biol, 38(8), 796–804. doi:10.1016/j.ibmb.2008.05.006

81. Loppin, B., Dubruille, R., & Horard, B. (2015). The intimate genetics of *Drosophila* fertilization. Open biology, 5(8), 150076.

82. Love, M. I., Huber, W., & Anders, S. (2014). Moderated estimation of fold change and dispersion for RNA-seq data with DESeq2. Genome Biol, 15(12), 550.

83. Lung, O., Kuo, L., & Wolfner, M. F. (2001). *Drosophila* males transfer antibacterial proteins from their accessory gland and ejaculatory duct to their mates. Journal of Insect Physiology, 47(6), 617–622.

84. Lung, O., & Wolfner, M. F. (2001). Identification and characterization of the major *Drosophila melanogaster* mating plug protein. Insect biochemistry and molecular biology, 31(6-7), 543–551.

85. Lv, D. K., Bai, X., Li, Y., Ding, X. D., Ge, Y., Cai, H., … Zhu, Y. M. (2010). Profiling of cold-stress-responsive miRNAs in rice by microarrays. Gene, 459(1-2), 39–47. doi:DOI 10.1016/j.gene.2010.03.011

86. Ma, W. C., Denlinger, D. L., Jarlfors, U., & Smith, D. S. (1975). Structural modulations in the tsetse fly milk gland during a pregnancy cycle. Tissue Cell, 7(2), 319–330. doi:0040-8166(75)90008-7 [pii]

87. Malik, A., Khan, K. M., Asif, M., Arsalan, H. M., Zahid, S., Manan, A., & Rasool, M. (2016). Development of Resistance Mechanism in Mosquitoes: Cytochrome P450, the Ultimate Detoxifier. Journal of Applied and Emerging Sciences, 4(2), pp100–117.

88. Marchini, D., Bernini, L. F., Marri, L., Giordano, P. C., & Dallai, R. (1991). The female reproductive accessory glands of the medfly *Ceratitis capitata*: antibacterial activity of the secretion fluid. Insect Biochemistry, 21(6), 597–605.

89. Masci, V. L., Di Luca, M., Gambellini, G., Taddei, A. R., Belardinelli, M. C., Guerra, L., … Fausto, A. M. (2015). Reproductive biology in Anophelinae mosquitoes (Diptera, Culicidae): fine structure of the female accessory gland. Arthropod structure & development, 44(4), 378–387.

90. McGraw, L. A., Clark, A. G., & Wolfner, M. F. (2008). Post-mating gene expression profiles of female *Drosophila melanogaster* in response to time and to four male accessory gland proteins. Genetics, 179(3), 1395–1408. doi:10.1534/genetics.108.086934

91. McKenna, D. D., Scully, E. D., Pauchet, Y., Hoover, K., Kirsch, R., Geib, S. M., … Arsala, D. (2016). Genome of the Asian longhorned beetle (*Anoplophora glabripennis*), a globally significant invasive species, reveals key functional and evolutionary innovations at the beetle–plant interface. Genome biology, 17(1), 227.

92. Meier, R., Kotrba, M., & Ferrar, P. (1999). Ovoviviparity and viviparity in the Diptera. Biological Reviews of the Cambridge Philosophical Society, 74(3), 199–258.

93. Michaud, M. R., Benoit, J. B., Lopez-Martinez, G., Elnitsky, M. A., Lee, R. E., & Denlinger, D. L. (2008). Metabolomics reveals unique and shared metabolic changes in response to heat shock, freezing and desiccation in the Antarctic midge, *Belgica antarctica*. Journal of insect physiology, 54(4), 645–655. doi:DOI 10.1016/j.jinsphys.2008.01.003

94. Mitchell, S. N., Kakani, E. G., South, A., Howell, P. I., Waterhouse, R. M., & Catteruccia, F. (2015). Evolution of sexual traits influencing vectorial capacity in anopheline mosquitoes. Science, 347(6225), 985–988.

95. Newfeld, S. J., Chartoff, E. H., Graff, J. M., Melton, D. A., & Gelbart, W. M. (1996). Mothers against dpp encodes a conserved cytoplasmic protein required in DPP/TGF-beta responsive cells. Development, 122(7), 2099–2108.

96. Olafson, P. U., Aksoy, S., Attardo, G. M., Buckmeier, G., Chen, X., Coates, C. J., … Friedrich, M. (2019). Functional genomics of the stable fly, *Stomoxys calcitrans*, reveals mechanisms underlying reproduction, host interactions, and novel targets for pest control. bioRxiv, 623009.

97. Orr-Weaver, T. L. (2015). When bigger is better: the role of polyploidy in organogenesis. Trends in Genetics, 31(6), 307–315.

98. Otti, O., McTighe, A. P., & Reinhardt, K. (2013). In vitro antimicrobial sperm protection by an ejaculate-like substance. Functional Ecology, 27(1), 219–226.

99. Otti, O., Naylor, R. A., Siva-Jothy, M. T., & Reinhardt, K. (2009). Bacteriolytic activity in the ejaculate of an insect. American Naturalist, 174(2), 292–295. doi:Doi 10.1086/600099

100. Palmer, W. J., & Jiggins, F. M. (2015). Comparative genomics reveals the origins and diversity of arthropod immune systems. Molecular biology and evolution, 32(8), 2111–2129.

101. Panfilio, K. A., Jentzsch, I. M. V., Benoit, J. B., Erezyilmaz, D., Suzuki, Y., Colella, S., … Ioannidis, P. (2019). Molecular evolutionary trends and feeding ecology diversification in the Hemiptera, anchored by the milkweed bug genome. Genome Biology, 20(1), 64.

102. Papa, F., Windbichler, N., Waterhouse, R. M., Cagnetti, A., D’Amato, R., Persampieri, T., … Papathanos, P. A. (2017). Rapid evolution of female-biased genes among four species of *Anopheles* malaria mosquitoes. Genome research, 27(9), 1536–1548.

103. Parisi, M., Nuttall, R., Edwards, P., Minor, J., Naiman, D., Lu, J., … Oliver, B. (2004). A survey of ovary -, testis-, and soma-biased gene expression in *Drosophila melanogaster* adults. Genome Biology, 5(6), R40. doi:10.1186/gb-2004-5-6-r40

104. Pauchet, Y., Wilkinson, P., Vogel, H., Nelson, D., Reynolds, S., Heckel, D. G., & Ffrench-Constant, R. (2010). Pyrosequencing the *Manduca sexta* larval midgut transcriptome: messages for digestion, detoxification and defence. Insect molecular biology, 19(1), 61–75.

105. Perry, J. C., Harrison, P. W., & Mank, J. E. (2014). The ontogeny and evolution of sex-biased gene expression in *Drosophila melanogaster*. Molecular biology and evolution, 31(5), 1206–1219.

106. Poiani, A. (2006). Complexity of seminal fluid: a review. Behavioral Ecology and Sociobiology, 60(3), 289–310.

107. Polak, M., Simmons, L. W., Benoit, J. B., Ruohonen, K., Simpson, S. J., & Solon-Biet, S. M. (2017). Nutritional geometry of paternal effects on embryo mortality. Proceedings of the Royal Society B: Biological Sciences, 284(1864), 20171492.

108. Price, C. S. (1997). Conspecific sperm precedence in *Drosophila*. Nature, 388(6643), 663.

109. Qazi, M. C. B., Heifetz, Y., & Wolfner, M. F. (2003). The developments between gametogenesis and fertilization: ovulation and female sperm storage in *Drosophila melanogaster*. Dev Biol, 256(2), 195–211.

110. Quinlan, M. E. (2016). Cytoplasmic streaming in the *Drosophila* oocyte. Annual review of cell and developmental biology, 32, 173–195.

111. Ramalho-Santos, J., Varum, S., Amaral, S., Mota, P. C., Sousa, A. P., & Amaral, A. (2009). Mitochondrial functionality in reproduction: from gonads and gametes to embryos and embryonic stem cells. Human reproduction update, 15(5), 553–572.

112. Rato, L., Alves, M. G., Socorro, S., Duarte, A. I., Cavaco, J. E., & Oliveira, P. F. (2012). Metabolic regulation is important for spermatogenesis. Nature Reviews Urology, 9(6), 330.

113. Raudvere, U., Kolberg, L., Kuzmin, I., Arak, T., Adler, P., Peterson, H., & Vilo, J. (2019). g: Profiler: a web server for functional enrichment analysis and conversions of gene lists (2019 update). Nucleic acids research.

114. Ravi Ram, K., & Wolfner, M. F. (2007). Seminal influences: *Drosophila* Acps and the molecular interplay between males and females during reproduction. Integrative and comparative biology, 47(3), 427–445.

115. Rial, D., Santos-Echeandía, J., Álvarez-Salgado, X. A., Jordi, A., Tovar-Sánchez, A., & Bellas, J. (2016). Toxicity of seabird guano to sea urchin embryos and interaction with Cu and Pb. Chemosphere, 145, 384–393.

116. Richard, M., & Hoch, M. (2015). *Drosophila* eye size is determined by innexin 2-dependent decapentaplegic signalling. Developmental Biology, 408(1), 26–40.

117. Rinehart, J. P., Hayward, S. A. L., Elnitsky, M. A., Sandro, L. H., Lee, R. E., & Denlinger, D. L. (2006). Continuous up-regulation of heat shock proteins in larvae, but not adults, of a polar insect. Proceedings of the National Academy of Sciences of the United States of America, 103(38), 14223–14227. doi:DOI 10.1073/pnas.0606840103

118. Rogers, D. W., Whitten, M. M., Thailayil, J., Soichot, J., Levashina, E. A., & Catteruccia, F. (2008). Molecular and cellular components of the mating machinery in *Anopheles gambiae* females. Proceedings of the National Academy of Sciences, 105(49), 19390–19395.

119. Ronges, D., Walsh, J. P., Sinclair, B. J., & Stillman, J. H. (2012). Changes in extreme cold tolerance, membrane composition and cardiac transcriptome during the first day of thermal acclimation in the porcelain crab *Petrolisthes cinctipes*. Journal of Experimental Biology, 215(Pt 11), 1824–1836. doi:10.1242/jeb.069658

120. Rosa, E., & Saastamoinen, M. (2017). Sex-dependent effects of larval food stress on adult performance under semi-natural conditions: only a matter of size? Oecologia, 184(3), 633–642.

121. Rosendale, A. J., Dunlevy, M. E., McCue, M. D., & Benoit, J. B. (2019). Progressive behavioural, physiological and transcriptomic shifts over the course of prolonged starvation in ticks. Molecular Ecology, 28(1), 49–65.

122. Rosendale, A. J., Romick-Rosendale, L. E., Watanabe, M., Dunlevy, M. E., & Benoit, J. B. (2016). Mechanistic underpinnings of dehydration stress in ticks revealed through RNA-seq and metabolomics. *Journal of Experimental Biology*, Submitted.

123. Rosetto, M., Belardinelli, M., Fausto, A., Marchini, D., Bongiorno, G., Maroli, M., & Mazzini, M. (2003). A mammalian-like lipase gene is expressed in the female reproductive accessory glands of the sand fly *Phlebotomus papatasi* (Diptera, Psychodidae). Insect molecular biology, 12(5), 501–508.

124. Sato, M., & Sato, K. (2011). Degradation of paternal mitochondria by fertilization-triggered autophagy in *C. elegans* embryos. Science, 334(6059), 1141–1144.

125. Scolari, F., Benoit, J. B., Michalkova, V., Aksoy, E., Takac, P., Abd-Alla, A. M., … Attardo, G. M. (2016). The spermatophore in *Glossina morsitans morsitans*: insights into male montributions to reproduction. Scientific reports, 6, 20334.

126. Shim, J., Gururaja-Rao, S., & Banerjee, U. (2013). Nutritional regulation of stem and progenitor cells in *Drosophila*. Development, 140(23), 4647–4656.

127. Shutt, B., Stables, L., Aboagye-Antwi, F., Moran, J., & Tripet, F. (2010). Male accessory gland proteins induce female monogamy in anopheline mosquitoes. Medical and veterinary entomology, 24(1), 91–94.

128. Sibley, P. K., Ankley, G. T., & Benoit, D. A. (2001). Factors affecting reproduction and the importance of adult size on reproductive output of the midge *Chironomus tentans*. Environmental Toxicology and Chemistry: An International Journal, 20(6), 1296–1303.

129. Silva, J. V., Yoon, S., Domingues, S., Guimarães, S., Goltsev, A. V., e Silva, E. F. d. C., … Fardilha, M. (2015). Amyloid precursor protein interaction network in human testis: sentinel proteins for male reproduction. BMC bioinformatics, 16(1), 12.

130. Sirot, L. K., Wong, A., Chapman, T., & Wolfner, M. F. (2015). Sexual conflict and seminal fluid proteins: a dynamic landscape of sexual interactions. Cold Spring Harbor perspectives in biology, 7(2), a017533.

131. Soller, M., Bownes, M., & Kubli, E. (1999). Control of oocyte maturation in sexually mature *Drosophila* females. Developmental biology, 208(2), 337–351. doi:10.1006/dbio.1999.9210

132. Sosic, D., & Olson, E. N. (2003). A new twist on twist: Modulation of the NF-KappaB pathway. Cell Cycle, 2(2), 75–77.

133. Strong, J. (1967). Ecology of terrestrial arthropods at Palmer station, Antarctic Peninsula. Entomology of Antarctica, 357–371.

134. Stroumbakis, N. D., Li, Z., & Tolias, P. P. (1996). A homolog of human transcription factor NF-X1 encoded by the *Drosophila* shuttle craft gene is required in the embryonic central nervous system. Molecular and cellular biology, 16(1), 192–201.

135. Sugg, P., Edwards, J. S., & Baust, J. (1983). Phenology and life history of *Belgica antarctica*, an Antarctic midge (Diptera: Chironomidae). Ecological Entomology, 8(1), 105–113.

136. Supek, F., Bošnjak, M., Škunca, N., & Šmuc, T. (2011). REVIGO summarizes and visualizes long lists of gene ontology terms. PLoS One, 6(7), e21800.

137. Swanson, W. J., Wong, A., Wolfner, M. F., & Aquadro, C. F. (2004). Evolutionary expressed sequence tag analysis of *Drosophila* female reproductive tracts identifies genes subjected to positive selection. Genetics, 168(3), 1457–1465.

138. Swevers, L., Raikhel, A., Sappington, T., Shirk, P., & Iatrou, K. (2005). Vitellogenesis and post-vitellogenic maturation of the insect ovarian follicle. In L.I. Gilbert, S. Gill, & K. Iatrou (Eds.), Comprehensive molecular insect science (pp. 87–155). Elsevier, NY

139. Teets, N. M., Elnitsky, M. A., Benoit, J. B., Lopez-Martinez, G., Denlinger, D. L., & Lee, R. E. (2008). Rapid cold-hardening in larvae of the Antarctic midge *Belgica antarctica*: cellular cold-sensing and a role for calcium. American Journal of Physiology-Regulatory Integrative and Comparative Physiology, 294(6), R1938–R1946. doi:DOI 10.1152/ajpregu.00459.2007

140. Teets, N. M., Kawarasaki, Y., Lee, R. E., Jr., & Denlinger, D. L. (2013). Expression of genes involved in energy mobilization and osmoprotectant synthesis during thermal and dehydration stress in the Antarctic midge, Belgica antarctica. Journal of Comparative Physiology B, 183(2), 189–201. doi:10.1007/s00360-012-0707-2

141. Teets, N. M., Peyton, J. T., Colinet, H., Renault, D., Kelley, J. L., Kawarasaki, Y., … Denlinger, D. L. (2012). Gene expression changes governing extreme dehydration tolerance in an Antarctic insect. Proceedings of the National Academy of Sciences of the United States of America, 109(50), 20744–20749. doi:10.1073/pnas.1218661109

142. Telfer, W. H., & Kunkel, J. G. (1991). The function and evolution of insect storage hexamers. Annual Review of Entomology, 36(1), 205–228.

143. Terashima, J., & Bownes, M. (2005). A microarray analysis of genes involved in relating egg production to nutritional intake in *Drosophila melanogaster*. Cell Death Differ, 12(5), 429.

144. Thailayil, J., Magnusson, K., Godfray, H. C. J., Crisanti, A., & Catteruccia, F. (2011). Spermless males elicit large-scale female responses to mating in the malaria mosquito *Anopheles gambiae*. Proceedings of the National Academy of Sciences, 108(33), 13677–13681.

145. Tian, C.-B., Wei, D., Xiao, L.-F., Dou, W., Liu, H., & Wang, J.-J. (2017). Comparative transcriptome analysis of three *Bactrocera dorsalis* (Diptera: Tephritidae) organs to identify functional genes in the male accessory glands and ejaculatory duct. Florida Entomologist, 100(1), 42–51.

146. Tian, E., & Hagen, K. G. T. (2007). O-linked glycan expression during *Drosophila* development. Glycobiology, 17(8), 820–827.

147. Tirmarche, S., Kimura, S., Dubruille, R., Horard, B., & Loppin, B. (2016). Unlocking sperm chromatin at fertilization requires a dedicated egg thioredoxin in *Drosophila*. Nature communications, 7, 13539.

148. Tran, D. T., & Ten Hagen, K. G. (2013). Mucin-type O-glycosylation during development. Journal of Biological Chemistry, 288(10), 6921–6929.

149. Tran, D. T., Zhang, L., Zhang, Y., Tian, E., Earl, L. A., & Ten Hagen, K. G. (2012). Multiple members of the UDP-GalNAc: polypeptide N-acetylgalactosaminyltransferase family are essential for viability in *Drosophila*. Journal of Biological Chemistry, 287(8), 5243–5252.

150. Turnier, J. L., Brunner, H. I., Bennett, M., Aleed, A., Gulati, G., Haffey, W. D., … Witte, D. (2018). Discovery of SERPINA3 as a candidate urinary biomarker of lupus nephritis activity. Rheumatology, 58(2), 321–330.

151. Usher, M. B., & Edwards, M. (1984). A dipteran from south of the Antarctic Circle: *Belgica Antarctica* (Chironomidae) with a description of its larva. Biological Journal of the Linnean Society, 23(1), 19–31.

152. Vanderperre, B., Cermakova, K., Escoffier, J., Kaba, M., Bender, T., Nef, S., & Martinou, J.-C. (2016). MPC1-like is a placental mammal-specific mitochondrial pyruvate carrier subunit expressed in postmeiotic male germ cells. Journal of Biological Chemistry, 291(32), 16448–16461.

153. Venancio, T., Cristofoletti, P., Ferreira, C., Verjovski-Almeida, S., & Terra, W. (2009). The *Aedes aegypti* larval transcriptome: a comparative perspective with emphasis on trypsins and the domain structure of peritrophins. Insect molecular biology, 18(1), 33–44.

154. Villarreal, S. M., Pitcher, S., Helinski, M. E., Johnson, L., Wolfner, M. F., & Harrington, L. C. (2018). Male contributions during mating increase female survival in the disease vector mosquito *Aedes aegypti*. Journal of insect physiology, 108, 1–9.

155. Vogt, C., Belz, D., Galluba, S., Nowak, C., Oetken, M., & Oehlmann, J. (2007). Effects of cadmium and tributyltin on development and reproduction of the non-biting midge *Chironomus riparius* (Diptera)—baseline experiments for future multi-generation studies. Journal of Environmental Science and Health Part A, 42(1), 1–9.

156. Weirauch, M. T., & Hughes, T. (2011). A catalogue of eukaryotic transcription factor types, their evolutionary origin, and species distribution. In A handbook of transcription factors (pp. 25–73): Springer.

157. Weirauch, M. T., Yang, A., Albu, M., Cote, A. G., Montenegro-Montero, A., Drewe, P., … Cook, K. (2014). Determination and inference of eukaryotic transcription factor sequence specificity. Cell, 158(6), 1431–1443.

158. Wensler, R. J., & Rempel, J. (1962). The morphology of the male and female reproductive systems of the midge, *Chironomus plumosus* L. Canadian Journal of Zoology, 40(2), 199–229.

159. Xie, T., & Spradling, A. C. (1998). Decapentaplegic is essential for the maintenance and division of germline stem cells in the *Drosophila* ovary. Cell, 94(2), 251–260.

160. Zhang, J., Marshall, K. E., Westwood, J. T., Clark, M. S., & Sinclair, B. J. (2011). Divergent transcriptomic responses to repeated and single cold exposures in *Drosophila melanogaster*. Journal of Experimental Biology, 214(Pt 23), 4021–4029. doi:10.1242/jeb.059535

161. Zhang, L., Zhang, Y., & Ten Hagen, K. G. (2008). A mucin-type O-glycosyltransferase modulates cell adhesion during *Drosophila* development. Journal of Biological Chemistry, 283(49), 34076–34086.

